# *CDK12* Loss Promotes Prostate Cancer Development While Exposing Vulnerabilities to Paralog-Based Synthetic Lethality

**DOI:** 10.1101/2024.03.20.585990

**Authors:** Jean Ching-Yi Tien, Yu Chang, Yuping Zhang, Jonathan Chou, Yunhui Cheng, Xiaoju Wang, Jianzhang Yang, Rahul Mannan, Palak Shah, Xiao-Ming Wang, Abigail J. Todd, Sanjana Eyunni, Caleb Cheng, Ryan J. Rebernick, Lanbo Xiao, Yi Bao, James Neiswender, Rachel Brough, Stephen J. Pettitt, Xuhong Cao, Stephanie J. Miner, Licheng Zhou, Yi-Mi Wu, Estefania Labanca, Yuzhuo Wang, Abhijit Parolia, Marcin Cieslik, Dan R. Robinson, Zhen Wang, Felix Y. Feng, Christopher J. Lord, Ke Ding, Arul M. Chinnaiyan

## Abstract

Biallelic loss of cyclin-dependent kinase 12 (*CDK12*) defines a unique molecular subtype of metastatic castration-resistant prostate cancer (mCRPC). It remains unclear, however, whether *CDK12* loss *per se* is sufficient to drive prostate cancer development—either alone, or in the context of other genetic alterations—and whether *CDK12*-mutant tumors exhibit sensitivity to specific pharmacotherapies. Here, we demonstrate that tissue-specific *Cdk12* ablation is sufficient to induce preneoplastic lesions and robust T cell infiltration in the mouse prostate. Allograft-based CRISPR screening demonstrated that *Cdk12* loss is positively associated with *Trp53* inactivation but negatively associated with *Pten* inactivation—akin to what is observed in human mCRPC. Consistent with this, ablation of *Cdk12* in prostate organoids with concurrent *Trp53* loss promotes their proliferation and ability to form tumors in mice, while *Cdk12* knockout in the *Pten*-null prostate cancer mouse model abrogates tumor growth. Bigenic *Cdk12* and *Trp53* loss allografts represent a new syngeneic model for the study of androgen receptor (AR)-positive, luminal prostate cancer. Notably, *Cdk12/Trp53* loss prostate tumors are sensitive to immune checkpoint blockade. *Cdk12*-null organoids (either with or without *Trp53* co-ablation) and patient-derived xenografts from tumors with *CDK12* inactivation are highly sensitive to inhibition or degradation of its paralog kinase, CDK13. Together, these data identify *CDK12* as a *bona fide* tumor suppressor gene with impact on tumor progression and lends support to paralog-based synthetic lethality as a promising strategy for treating *CDK12-*mutant mCRPC.

## INTRODUCTION

Cyclin-dependent kinases (CDKs) fall into two major categories: cell cycle regulatory CDKs (CDK4 and 6), which drive progression through the cell cycle, and transcriptional CDKs (CDK7, 8, 9, 12 and 13), which regulate gene expression^1^. CDK12 is a transcriptional cyclin-dependent kinase that associates with DNA in protein-coding and enhancer regions^2^. Upon binding its cognate cyclin (cyclin K), CDK12 phosphorylates Ser2 residues within the C-terminal domain (CTD) of RNA polymerase II (Pol-II)^3^. This modification facilitates recruitment of phosphorylated CTD-associated proteins required for transcriptional elongation^3–5^—a process that has been suggested to be particularly important for expression of longer genes containing numerous exons^4^. CDK12 further controls gene expression via regulation of alternative splicing^6,7^ and pre-mRNA processing^8,9^. Notably, genetic and pharmacologic targeting of CDK12 in several systems show CDK12 to be a key transcriptional regulator of genes mediating DNA damage responses^4,10–12^, and *in vitro* CDK12 loss-of-function results in reduced RAD51 foci formation and PARP inhibitor sensitivity^4,13–15^.

CDK13 is a vertebrate paralog of CDK12 and shares 92% homology in the kinase domain and a similar three-dimensional structure^1^. In line with this, CDK13 also binds cyclin K to exert Pol-II CTD kinase activity^16^. Comparison of gene expression profiles from HCT116 colon cancer cells subjected to *CDK12* or *CDK13* knockdown demonstrated 75% overlap in affected transcripts^11^. Fan et al. applied CRISPR-Cas9 homology-directed repair to generate MV4-11 leukemia cell lines in which CDK12, CDK13, or both could be inhibited via administration of ATP analog NM-PPI^17^. This approach revealed that inhibition of both kinases, but not either individually, was sufficient to yield broad reduction in CTD phosphorylation, reduced proliferation, and cell death^17^. In both systems, transcripts related to DNA damage response were found to be predominantly regulated by CDK12^11^. Therefore, while maintenance of genomic stability depends on CDK12, maintenance of cell proliferation and survival is enabled via redundant actions of CDK12 and CDK13.

Biallelic inactivation and mutations of *CDK12* have been identified in several malignancies^18–20^. For instance, biallelic *CDK12* loss is observed in about 4% of serous ovarian carcinoma, characterizing a disease subtype in which genomic instability is driven by recurrent focal tandem duplications^20^. Notably, these tumors are genetically distinct from those bearing inactivating mutations in the homologous recombination regulators *BRCA1* and *BRCA2*^20^. Whole-exome sequencing of a cohort of 360 metastatic castration-resistant prostate cancer (mCRPC) samples from Stand Up to Cancer (SU2C)^21^, Mi-Oncoseq^22^, and University of Michigan rapid autopsy cases^23^ revealed biallelic alterations in 25/360 of mCRPC cases (6.9%). By contrast, sequencing of primary prostate cancers in The Cancer Genome Atas (TCGA) revealed only 6/498 cases (1.2%) harboring biallelic *CDK12* alterations, suggesting that *CDK12* mutations are markedly enriched in metastatic disease^18^. Biallelic loss of *CDK12* constituted a unique mCRPC subtype, genetically distinct from those driven by other primary driver genes—including ETS gene fusions, *SPOP* mutations, homologous recombination deficiency (HRD), and mismatch repair deficiency (MMRD)^21–27^. *CDK12*-mutant tumors were characterized by a genomic instability pattern like that described in ovarian cancer^20^, in which recurrent gains secondary to focal tandem duplications yielded putative neo-antigens^18^. To this end, *CDK12* loss was associated with T cell infiltration of mCRPC tumors^18^. Despite these associations, it remains unclear whether *CDK12* represents a *bona fide* tumor suppressor gene which, when inactivated, can drive tumorigenesis. Furthermore, it is unclear whether *CDK12* loss renders prostate tumors susceptible to paralog-based synthetic lethality^28^.

Here, we developed *in vivo* and *in vitro* systems to test the impact of *Cdk12* ablation— both independently and in the context of other canonical mCRPC-related mutations identified in unbiased CRISPR screening. We provide compelling evidence that *Cdk12* is a tumor suppressor gene whose loss enhances tumorigenesis and progression in the setting *of Trp53* loss, but by contrast, inhibits tumor growth in the context of *Pten* loss. *Cdk12* loss in the mouse prostate induces robust T cell infiltration analogous to what is observed in human *CDK12*-mutant tumors^18^. We establish bigenic *Cdk12/Trp53* loss as a novel syngeneic model of prostate cancer that, unlike other prostate genetically engineered mouse models (GEMMs), exhibits an androgen receptor-positive (AR+) luminal phenotype. Furthermore, *Cdk12/Trp53* loss prostate tumors respond to immune checkpoint blockade. Finally, we leveraged paralog-based synthetic lethality to demonstrate that murine and human tumor tissue lacking functional CDK12 is particularly sensitive to CDK13 inhibition and degradation—a finding with clinical relevance for the treatment of *CDK12*-mutant cancers.

## RESULTS

### Prostate-specific CDK12 ablation induces preneoplastic lesions and T cell infiltration

As mentioned above, biallelic loss-of-function mutations in the *CDK12* gene occur in ∼7% of mCRPC^18^, but whether *CDK12* inactivation contributes to prostate tumorigenesis is unknown. To resolve this question, we employed genetically engineered mice in which exons 3 and 4 of the *Cdk12* gene are flanked by lox-P sites (*Cdk12^f/f^* mice)^29^. Cre-mediated recombination of these loci excises the CDK12 kinase domain to yield a nonfunctional, truncated protein product and reduced mRNA levels^10^. We crossed *Cdk12^f/f^* mice into the established probasin Cre (*Pb-Cre*) line^30^. The resulting mixed genetic background animals (*Cdk12^pc-/-^* mice) lack functional CDK12 in prostate epithelial cells **(Fig S1A)**.

In the *Pb-Cre* model, roughly half of the luminal epithelial cells (LECs) and a smaller proportion of basal cells (BCs) have Cre activity. Expression varies by lobe, such that active Cre is present in most luminal cells of the ventral prostate (VP), dorsal prostate (DP), and lateral (LP) prostate but is present in less than half of luminal cells in the anterior prostate (AP). In our system, we confirmed a similar pattern of CDK12 ablation, showing *Cdk12^pc-/-^* mice to have loss of CDK12 protein expression in 38%, 59%, 60%, and 70% of AP, VP, DP, and LP epithelial cells, respectively **(Fig S1B-C)**. *In situ* hybridization (ISH) broadly corroborated the IHC findings, showing similarly distributed *Cdk12* transcript loss **(Fig S1B)**.

Young *Cdk12^pc-/-^* mice displayed no significant changes in prostate histology (**Fig S1B)**. On the contrary, prostates of 52-week-old *Cdk12^pc-/-^* mice exhibited regions of epithelial hyperplasia with loss of nuclear polarity and isonucleosis. Histologically atypical tissue occupied roughly 2% cross-sectional area in the *Cdk12^pc-/-^* anterior, ventral, and lateral prostate, while accounting for 10% of cross-sectional area in the dorsal prostate. In contrast, similar tissue accounted for < 1% of cross-sectional area in all lobes of wild-type (WT) littermates **(Fig S2A-C)**. To mitigate genetic variability in the system, we generated pure background *Cdk12* knockout mice by backcrossing *Cdk12^pc-/-^* animals into the C57BL/6 background over six generations. The resulting animals showed more significant prostate atypia than the mixed background model—including areas of focal high-grade prostatic intraepithelial neoplasia (HGPIN) and atypical intraductal proliferation (AIP) **(Fig 1A-B)**. Pathological scoring showed these higher grade lesions to be completely absent in the WT prostate but to occupy approximately 5% of total prostate cross-sectional area in *Cdk12^pc-/-^* mice. Similarly, hyperplastic lesions occupied 12% of prostate cross-sections in *Cdk12^pc-/-^* mice but <5% in WT controls **(Fig 1B)**. Neoplastic lesions in the *Cdk12^pc-/-^* prostate were characterized by p63 basal cell accumulation **(Fig 1C)** and increased cellular proliferation as indicated by Ki67 immunopositivity **(Fig 1D).**

**Figure 1.**
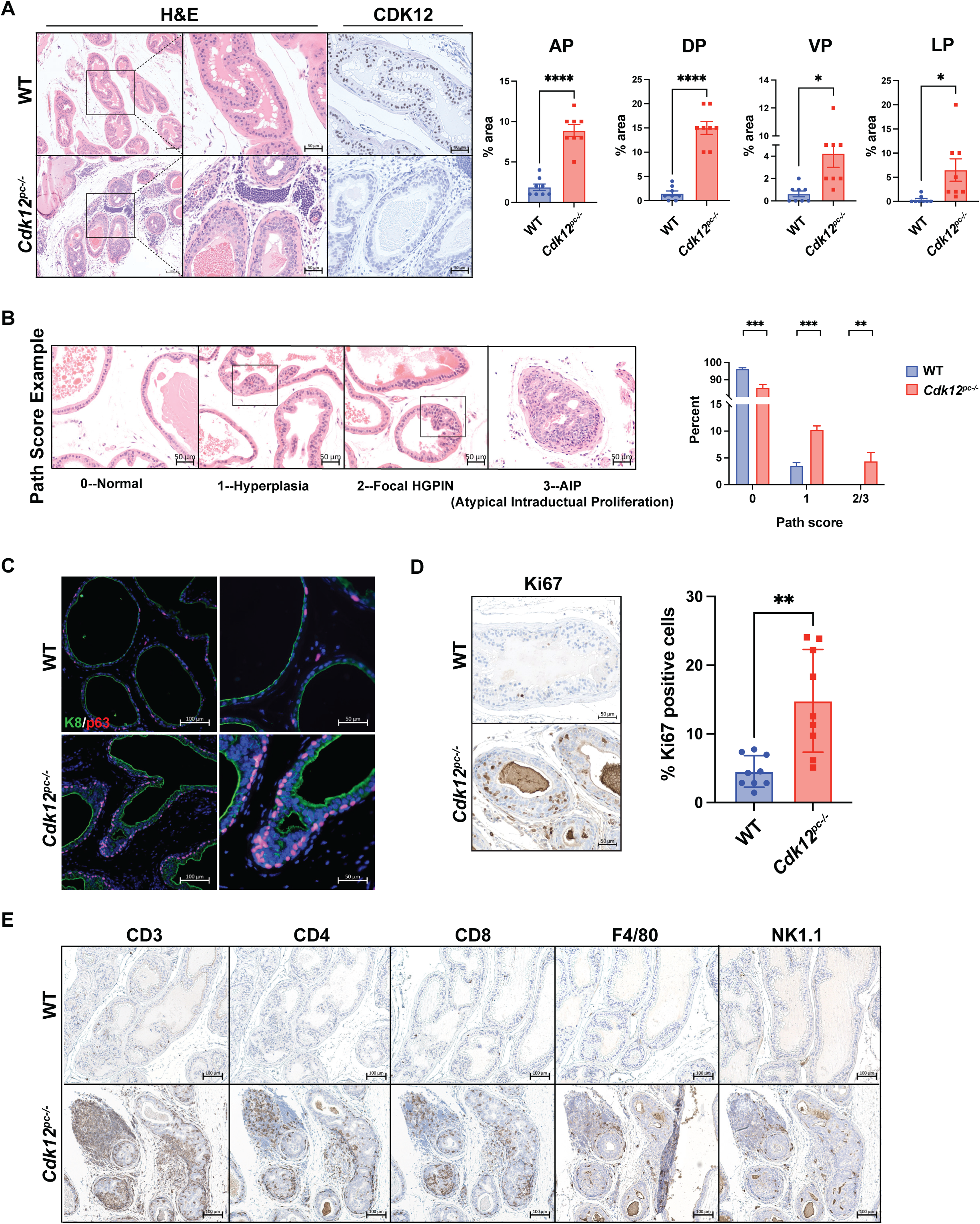
*Cdk12* ablation in the prostate epithelium induces neoplasia. **(A)** H&E staining and CDK12 immunohistochemistry (IHC) in representative prostate samples from 52-week-old *Cdk12^pc-/-^* mice (pure C57 background) and wild-type (WT) controls. Scale bars in left panel indicate 100μm. Other scale bars indicate 50μm.Bar graphs indicate percent cross-sectional area in each prostate lobe occupied by histologically abnormal tissue. Anterior prostate, AP. Dorsal prostate, DP. Ventral prostate, VP. Lateral prostate, LP. (n=8 mice per group). **(B)** Pathological scoring (Path score) of prostate tissue from the same animals. Numerical scores assigned to normal tissue (0), hyperplasia (1), focal high grade prostatic intraepithelial neoplasia (HGPIN) (2) and atypical intraductal proliferation (AIP) (3) (as indicated by respective images). Scale bars indicate 50μm. Bar graph shows percentage of prostate cross-sectional area occupied by tissue of each path score (scores 2 and 3 added together). (n=7-8 mice per group). **(C)** Immunofluorescent staining of cytokeratin-8 (K8) and p63. 52-week-old *Cdk12^pc-/-^* image shows an area of focal HGPIN with expansion of p63(+) basal cells. Scale bars in left and right images respectively represent 100μm and 50μm. **(D)** Ki67 immunohistochemistry from *Cdk12^pc-/-^* or WT mice. Scale bars indicate 50μm. Bar graph indicates percentage Ki67(+) cells per high powered field. (n=9 images from 3 mice per group). **(E)** Immunohistochemistry for immune cell markers, indicative of a T cell predominant infiltrate surrounding neoplastic lesions in *Cdk12^pc-/-^* animals. Scale bars indicate 100μm. Data are represented as mean ± SEM. t-test used individual comparisons, one way ANOVA for multiple comparisons, *p<0.05, **p<0.01, ***p<0.001, ****p<0.0001.

Human prostate cancers with biallelic *CDK12* inactivation develop T cell-predominant immune infiltrates^18^. A similar pattern emerged in the *Cdk12^pc-/-^*prostate, as regions surrounding lesions were enriched in CD3, CD4, and CD8(+) cells **(Fig 1E)**. Infiltration of F4/80(+) macrophages and NK1.1(+) natural killer cells was comparatively limited **(Fig 1E)**. In summary, *Cdk12* loss *per se* is sufficient to induce widespread preneoplastic changes with corresponding immune infiltrates in the mouse prostate.

### Cdk12-null prostate-derived organoids are hyperplastic and display basal-luminal disorganization

To better define the tumorigenic potential brought about by *Cdk12* loss, we generated organoids from pure populations of *Cdk12*-null prostate epithelial cells. Given the mixture of CDK12(+) and CDK12(-) cells in the *Cdk12^pc-/-^* prostate epithelium **(Fig S1B)**, doing so required a system in which cells with *Cdk12* ablation could be identified and isolated. We, therefore, intercrossed *Cdk12^pc-/-^* mice with *mT/mG* reporter mice. The latter animals harbor a constitutively expressed *td-Tomato/ stop-floxed eGFP* construct. At baseline, *mT/mG* cells express *td-Tomato* [Tom(+)] and appear red. In the setting of active Cre recombinase, the *td-Tomato* construct is excised and *eGFP* expressed [GFP(+)], such that cells appear green **(Fig S3A)**. In *Pb-Cre*;*Cdk12^f/f^*;*mT/mG* mice, cells with active probasin Cre (and resultant *Cdk12* ablation) are GFP(+)/ green.

We used flow cytometry to isolate GFP(+) and Tom(+) basal and luminal epithelial cells from prostates of 52-week-old *Pb-Cre*;*Cdk12^f/f^*;*mT/mG* mice **(Fig S3B)** and observed the expected reduction of *Cdk12* transcript in GFP(+) cells **(Fig S3C)**. GFP(+) and Tom(+) basal cells respectively gave rise to organoids that were uniformly green and red in color **(Fig S3D)**. Since luminal epithelial cell-derived organoids lacked this strict color segregation **(Fig S3B)**, we employed only basal cell-derived organoids for subsequent experiments. Upon initial analysis, red *Cdk12^WT^* organoids exhibited normal murine prostate morphology—characterized by an organized epithelial layer surrounding a large lumen^31^. By contrast, green *Cdk12^KO^* organoids were smaller in size and lacked lumens **(Fig 2A, Fig S3D)**.

**Figure 2.**
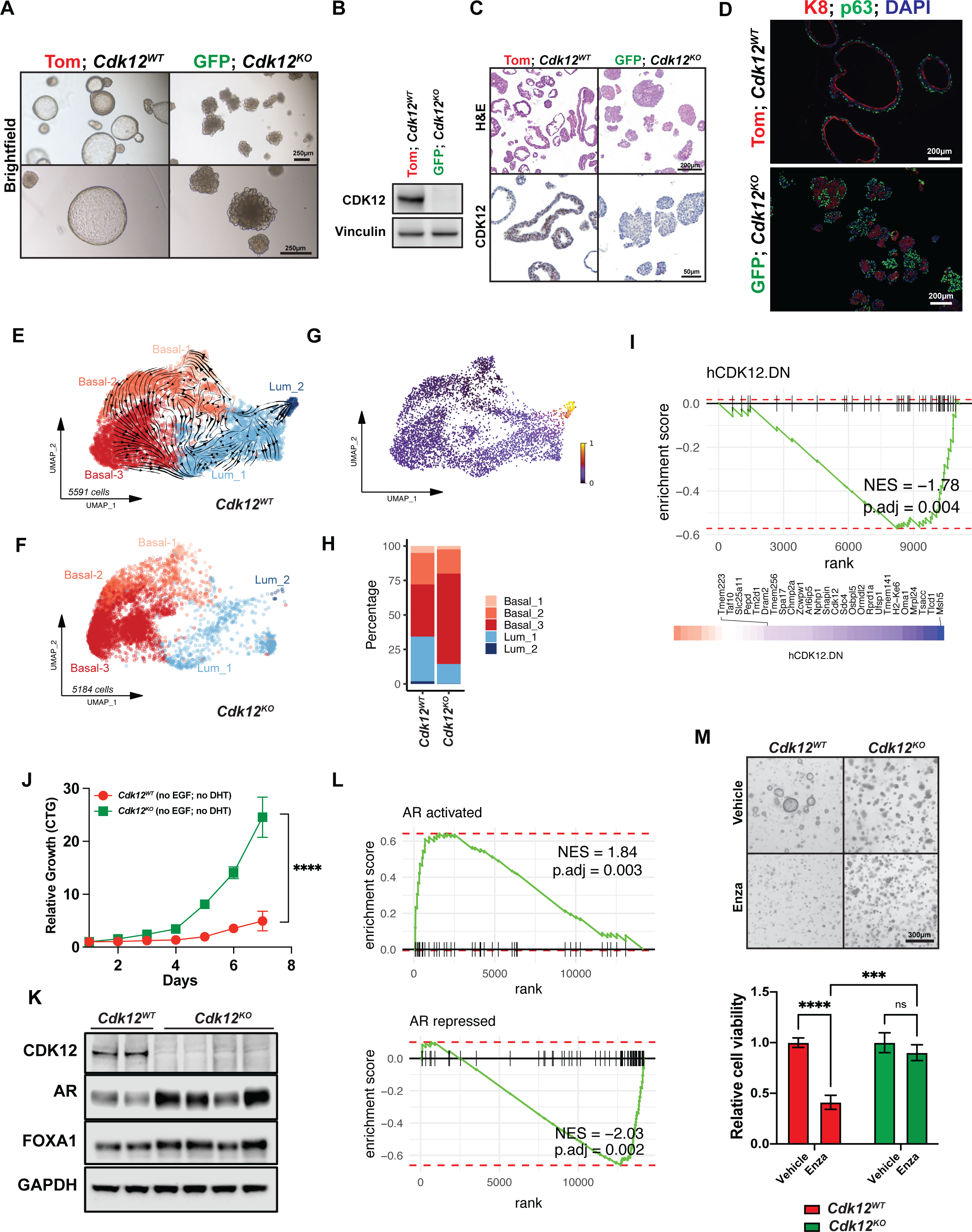
Organoids derived from the *Cdk12^pc-/-^* prostate are morphologically abnormal, with impaired basal-luminal segregation and reduced androgen dependence. **(A)** Brightfield images of organoids derived from *Pb-Cre*;*Cdk12^f/f^*;*mT/mG* prostate basal cells (52-week time point). Tom indicates Td-tomato-expressing cells with wild-type *Cdk12* (*Cdk12^WT^*). GFP indicates GFP-expressing cells with *Cdk12* ablation (*Cdk12^KO^*). Scale bars indicate 200μm. **(B)** Western blot analysis of organoids from each group demonstrating CDK12 loss. Vinculin serves as a loading control. **(C)** *Cdk12^KO^* organoid morphology: H&E staining and CDK12 immunohistochemistry. Scale bars in top and bottom panels respectively indicate 200μm and 100μm. **(D)** Immunofluorescence for cytokeratin-8 (K8) and p63 indicates basal-luminal disorganization in *Cdk12^KO^* organoids. Scale bars indicate 200μm. **(E)** UMAP of scRNA-seq from *Cdk12^WT^* organoids (derived from 3 mice). The streamlines with arrows obtained from RNA velocity analysis indicate directions of cell progression from one state to another. **(F)** Cells from *Cdk12^KO^* organoids colored by pseudo-time; the five identified cell states progress from Basal_1, Basal_2, Basal_3, Lum_1, to Lum_2. **(G)** Cells from *Cdk12^KO^* organoids (derived from 3 mice) projected into the UMAP of *Cdk12^WT^*. Pseudocolor indicates presence (yellow) or absence (purple) of *Cdk12* transcript. **(H)** Distributions of different cell states in *Cdk12^WT^* and *Cdk12^KO^*organoids. *Cdk12^KO^* organoids are less differentiated with increased basal cell accumulation. The most differentiated (Lum_2) population is lost in *Cdk12^KO^* organoids. **(I)** Enrichment in *Cdk12^KO^* organoids of down-regulated genes from human prostate cancer with *CDK12* inactivation [as defined in^18^]. **(J)** Proliferation of *Cdk12^WT^* and *Cdk12^KO^* organoids grown in the absence of epidermal growth factor (EGF) and dihydrotestosterone (DHT) as measured by the Cell Titer-Glo (CTG) assay. (n= 3 replicates per group in 2 unique experiments). **(K)** Protein expression of CDK12, AR, and FOXA1 in multiple monoclonal *Cdk12^WT^* and *Cdk12^KO^* organoid lines. GAPDH serves as a loading control. **(L)** Gene set enrichment of AR target genes (activated and repressed) in *Cdk12^WT^* and *Cdk12^KO^* organoids. **(M)** Morphology and viability quantification of *Cdk12^WT^* and *Cdk12^KO^*organoids subjected to enzalutamide (Enza) treatment. (n= 3 replicates per group in 2 unique experiments) Data are represented as mean ± SEM. Two-way ANOVA used for statistical comparisons, ***p<0.001; ****p<0.0001; ns, not significant.

We applied multiple experimental systems to confirm that this phenotype resulted specifically from *Cdk12* loss. First, we compared GFP(+) basal cell-derived organoids from *PbCre*;*Cdk12^f/f^*;*mT/mG* mice and *PbCre*;*Cdk12^+/+^*;*mT/mG* mice. Organoids derived from the former (*Cdk12*-null) were small and lacked lumens, whereas those derived from the latter (*Cdk12*-intact) had normal morphology **(Fig S3E)**. Next, we isolated basal cells from *Cdk12^f/f^*;*mT/mG* mice and then treated *in vitro* with either Cre-expressing adenovirus or control adenovirus before generating organoids. Only organoids from Cre-expressing adenovirus-treated (*Cdk12*-null) cells demonstrated size reduction and absent lumens **(Fig S3F)**.

We, therefore, conducted further experiments comparing red *Cdk12^WT^*and green *Cdk12^KO^* organoids **(Fig S3D)**. In *Cdk12^KO^* organoids, we confirmed loss of CDK12 protein with western blot **(Fig 2B)** and immunohistochemistry **(Fig 2C)**. Detailed observation of their morphology revealed *Cdk12^KO^* organoids to be hyperplastic with disorganization of K8(+) luminal epithelial cells and p63(+) basal cells **(Fig 2D)**. Single cell RNA sequencing (scRNA-seq) (with velocity analysis) confirmed this, demonstrating reduced basal to luminal cell differentiation in *Cdk12^KO^*versus *Cdk12^WT^* organoids **(Fig 2E-G)**, leading to basal cell accumulation in *Cdk12^KO^* samples **(Fig 2H)**. Strikingly, pseudo-bulk analysis showed that gene sets altered in human prostate cancer with biallelic *CDK12* loss were significantly enriched in *Cdk12^KO^* organoids **(Fig 2I)**.

Given the relationship between *CDK12* loss and castration resistance in human prostate cancer, we evaluated growth of *Cdk12^KO^* organoids in testosterone-depleted culture medium. We observed that *Cdk12^KO^* organoids demonstrated increased proliferation over *Cdk12^WT^* organoids in this setting **(Fig 2J),** while also exhibiting increased protein levels of AR and the AR target gene FOXA1 **(Fig 2K)**. *Cdk12^KO^* organoids also exhibited enrichment of gene expression signatures consistent with increased AR signaling **(Fig 2L)**. To this end, *Cdk12* loss also rendered organoids resistant to the anti-androgen enzalutamide, whereas *Cdk12^WT^* organoids displayed significantly decreased cell viability with enzalutamide treatment **(Fig 2M)**. Interestingly, *CDK12*-mutant prostate cancer patients display poorer responses to hormonal therapies as compared to *CDK12* wild-type prostate cancers^32,33^. Taken together, these data demonstrate that *Cdk12^KO^* organoids display an abnormal phenotype consistent both with the neoplastic lesions seen in prostates of *Cdk12^pc-/-^* mice and with *CDK12*-mutant human prostate cancer.

### In vivo CRISPR screen demonstrates Cdk12 loss is positively associated with p53 inactivation

Clinically, *CDK12* inactivation displays variable overlap with other cardinal prostate cancer mutations^18^, suggesting the impact of *CDK12* loss on tumorigenesis is related to mutational context. To define the mutations that positively and negatively interact with *Cdk12* loss, we applied CRISPR screening in our organoid model. Specifically, we employed a MusCK library that contains five sgRNAs per gene and over 4500 genes that are implicated in tumor initiation, progression, and immune modulation^34^. We transduced the library into *Cdk12^KO^* organoids with MOI <0.3. After puromycin selection, we transplanted library-transduced organoids subcutaneously into immunocompromised NSG mice. After six months, we harvested the resulting tumors and sequenced to compare sgRNA abundance versus freshly infected organoids prior to *in vivo* transplantation **(Fig 3A)**. Strikingly, *Trp53*, a gene commonly inactivated in *CDK12*-mutant human tumors^18^, emerged as the most significantly depleted gene in the screen (i.e., its sgRNAs were most abundant) **(Fig 3B)**.

**Figure 3.**
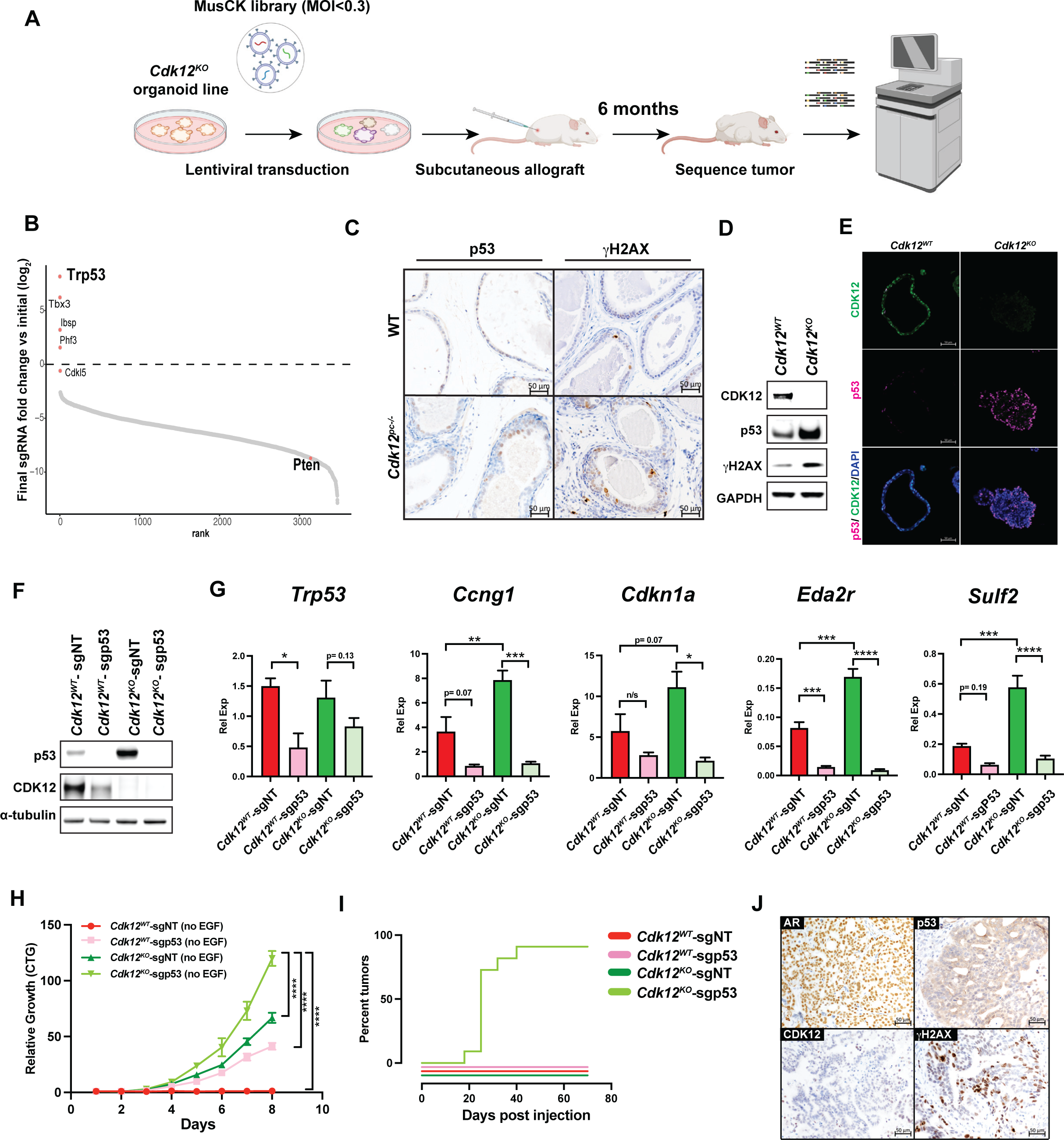
*Cdk12* and *Trp53* inactivating mutations interact to promote prostate cancer. **(A)** Workflow for CRISPR library screening of *Cdk12* interacting genes. *Cdk12^KO^*organoid cells transduced with MusCK CRISPR library at multiplicity of infection (MOI) 0.3. Tumor allografts generated from transfected cells grown subcutaneously in immunocompromised mice for six months. Tumors harvested and subjected to sequencing analysis to determine enrichment or depletion of library guide RNAs. **(B)** Snake plot representing log2 fold change of guide RNAs in sequenced tumor samples described in (A). **(C)** Immunohistochemistry for p53 (left panels) and DNA damage marker (γH2AX; right panels) in prostates of one-year-old WT and *Cdk12^pc-/-^* mice. Scale bars indicate 50μm. **(D)** Protein expression of p53 and γH2AX in *Cdk12^WT^* and *Cdk12 ^KO^* organoids. GAPDH serves as a loading control. **(E)** CDK12-p53 co-staining in *Cdk12^WT^* and *Cdk12^KO^* organoids. Scale bars indicate 50μm. **(F)** CRISPR-mediated *Trp53* ablation in *Cdk12^WT^* and *Cdk12^KO^* organoids. sgp53 indicates *Trp53*-specific guide RNA. sgNT indicates control non-targeting guide RNA. Alpha tubulin serves as a loading control. **(G)** Relative expression (Rel Exp) levels of *Trp53* and p53 target genes in samples described in (F). (n= 3 samples per group). **(H)** Cell proliferation in organoids from groups indicated in (F) as measured by CTG assay. (n= 3-4 samples per group) **(I)** Kaplan Meier plots indicating tumor formation during the 70 days post implantation (subcutaneous allograft) in *Cdk12^WT^* and *Cdk12^KO^* organoids with or without *Trp53* ablation. Note that only *Cdk12^KO^*-sgp53 cells form viable allograft tumors. (n= 5 mice each with 2 tumor injection sites per group). **(J)** Immunohistochemical staining of AR, p53, CDK12, and γH2AX in *Cdk12^KO^*-sgp53 allografts. Scale bars indicate 50μm. Data are represented as mean ± SEM. Two-way ANOVA was used for statistical comparisons. *p<0.05; **p<0.01; ***p<0.001; ****p<0.0001.

Notably, in prostates of *Cdk12^pc-/-^* mice, we observed both increased DNA damage (as indicated by γH2AX immunohistochemistry) and consequent p53 protein induction within preneoplastic lesions **(Fig 3C)**. In agreement, p53 signaling was among the most significantly upregulated pathways on scRNA-seq analysis of luminal cells isolated from the *Cdk12*-null prostate **(Fig S4A-E)**. *Cdk12^KO^* organoids phenocopied the *in vivo* findings, exhibiting elevated abundance of p53 and γH2AX protein **(Fig 3D-E)**. In summary, *Cdk12* loss induces DNA damage and p53 expression, while cells with concomitant *Trp53* loss are preferentially enriched in allografts derived from *Cdk12^KO^* organoids.

### Concomitant Trp53 loss enhances tumorigenic potential of Cdk12-null prostate epithelial cells

We hypothesized that p53 induction in *Cdk12*-null prostate epithelial cells represented a response to increased DNA damage, and hence, that these cells would display enhanced tumorigenic potential in the setting of *Trp53* loss. We tested this hypothesis by using CRISPR-Cas9 to ablate the *Trp53* gene in *Cdk12^WT^* and *Cdk12^KO^* organoids. We observed that both p53 protein levels **(Fig 3F)** and target gene expression **(Fig 3G)** were higher in *Cdk12^KO^* organoids than in *Cdk12^WT^* organoids when each was transfected with control sgRNA (*Cdk12^KO^*-sgNT and *Cdk12^WT^*-sgNT, respectively). Transfection of sgp53—to generate *Cdk12^WT^* -sgp53 and *Cdk12^KO^*-sgp53 organoids—yielded the appropriate reductions in p53 protein and target gene expression **(Fig 3F-G)**. Organoids lacking either *Trp53* or *Cdk12* alone (*Cdk12^WT^*-sgp53 and *Cdk12^KO^*-sgNT organoids, respectively) proliferated more rapidly than pure WT (*Cdk12^WT^*-sgNT) organoids. Strikingly, organoids lacking both genes (*Cdk12^KO^*-sgp53 organoids) proliferated more rapidly than either single-knockout organoid type **(Fig 3H)**.

We then implanted organoid cells as subcutaneous allografts in immunocompromised mice. While WT-sgNT, WT-sgp53, and *Cdk12^KO^*-sgNT organoids failed to form tumors during a 60-day monitoring period, 100% of *Cdk12^KO^*-sgp53 organoids generated tumors within 50 days of implantation **(Fig 3I)**. Immunohistochemistry for AR, p53, and γH2AX showed resulting tumors to be of prostate origin and to display DNA damage **(Fig 3J)**. Furthermore, tumors could be serially passaged in mice, exhibiting improved growth with each passage **(Fig S5A)**. Together, these data reveal that concomitant loss of *Cdk12* and *Trp53*—respectively responsible for DNA maintenance and DNA damage responses—drives tumorigenesis beyond the loss of either factor alone. These findings are in line with clinical data that demonstrate frequent association between inactivating mutations in *CDK12* and *TP53* in mCRPC^18^.

### Cdk12/Trp53 double knockout allografts exhibit lymphocytic immune responses and increased sensitivity to immune checkpoint inhibitor therapy

To assess the immunogenicity of *Cdk12^KO^*-sgp53 organoid-derived tumors, we developed a novel syngeneic allograft line by implanting them subcutaneously in immunocompetent C57BL/6 mice. In this system, they maintained their AR(+), K8(+), p63(+) histology while demonstrating pronounced γH2AX staining **(Fig S5B)**. Despite growing similarly to established prostate cancer models **(Fig 4A),** *Cdk12^KO^*-sgp53 allografts elicited a T cell-predominant immune infiltrate that was not observed in Myc-CaP or TRAMP-C2 allografts or prostate tumors of *Pten*-null mice **(Fig 4B)**. Most notably, CD8(+) T cells broadly permeated *Cdk12^KO^*-sgp53 allografts but were essentially absent in Myc-CaP, TRAMP-C2, and *Pten^pc-/-^* samples.

**Figure 4.**
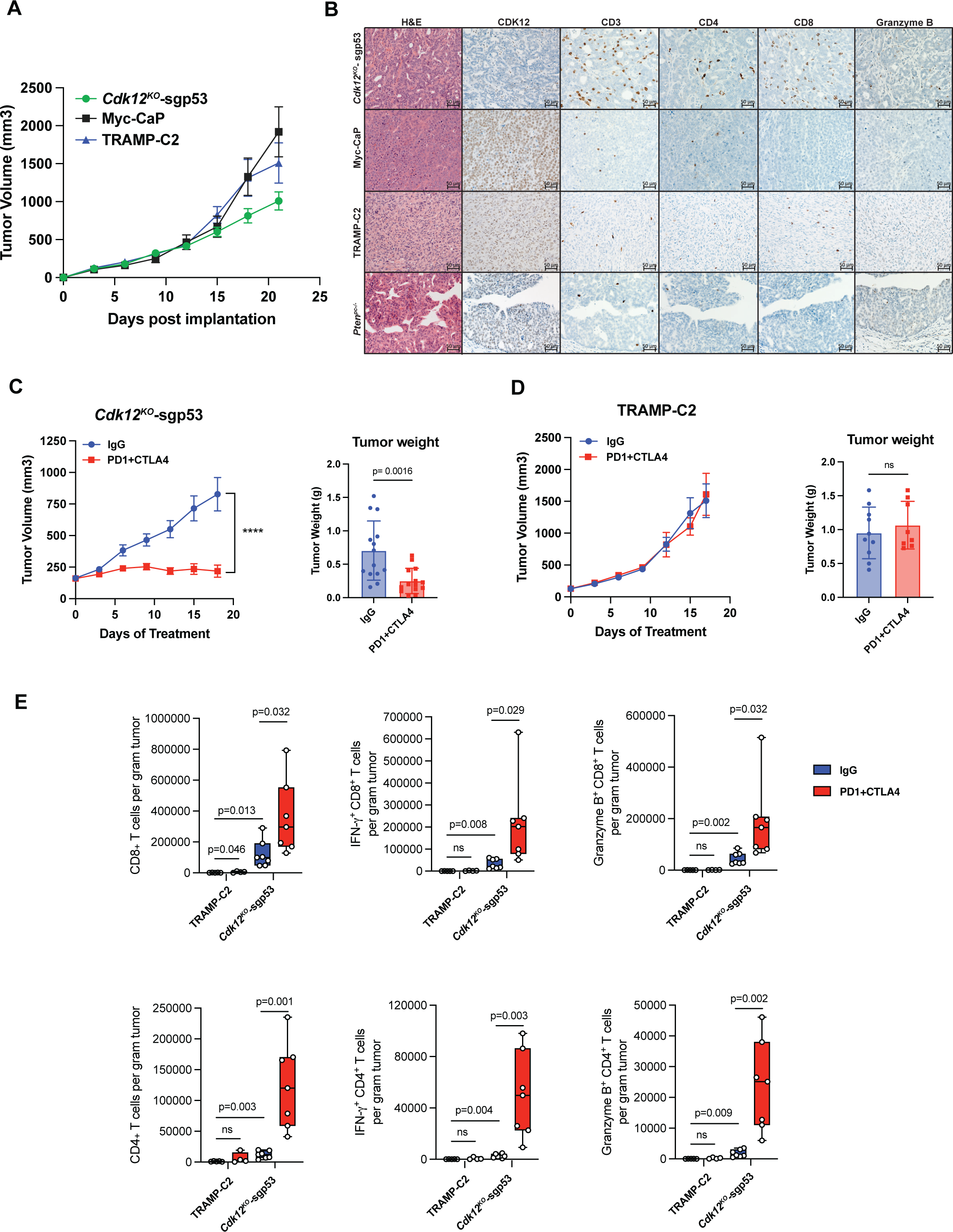
*Cdk12/Trp53* double knockout allografts exhibit lymphocytic immune responses and increased sensitivity to immune checkpoint inhibitor therapy. **(A)** Growth of *Cdk12^KO^*-sgp53, Myc-CaP, and TRAMP-C2 allografts in immunocompetent wild type mice. (n=10-15 mice, each with 2 tumor injection sites, per group). **(B)** Immunohistochemical staining of CDK12, T cell markers (CD3, CD4, CD8), and NK cell marker granzyme B in *Cdk12^KO^*-sgp53 allografts, Myc-CaP allografts, and TRAMP-C2 allografts, and prostates of the established *Pten^pc-/-^* prostate cancer mouse model. Scale bars indicate 50μm. **(C-D)** Tumor volume time course and endpoint weights of *Cdk12^KO^*-sgp53 (C) and TRAMP-C2 **(D)** allografts subjected to treatment with anti-PD1/CTLA4 cocktail. (n= 7-8 mice, each with 2 tumor injection sites, per group). (**E**) Flow cytometry-based quantification of CD4(+) and CD8(+) T cells (total, IFNγ(+), granzyme B(+)) in *Cdk12^KO^*-sgp53 and TRAMP-C2 allograft samples +/- treatment with anti-PD1/ CTLA4 cocktail. (n= 7-8 samples per group). Data are represented as mean ± SEM. Two-way ANOVA test was used for statistical comparison of tumor volume for (C) and in (E) and unpaired t test was used for tumor weight in (C) and (D). ****p<0.0001; ns, not significant.

Given these findings, we hypothesized that *Cdk12^KO^*-sgp53 allografts would be sensitive to immune checkpoint inhibitors. Indeed, equivalent doses of an anti-PD1/CTLA4 antibody cocktail strongly inhibited the growth and weight of these tumors **(Fig 4C)** but failed to curb that of TRAMP-C2 allografts **(Fig 4D)**. Strikingly, immune profiling of both tumor types revealed significantly more CD4(+) and CD8(+) T cells in *Cdk12^KO^*-sgp53 versus TRAMP-C2 samples **(Fig 4E)**. These differences were particularly pronounced in the setting of anti-PD1/CTLA4 therapy.

Interestingly, heightened immunogenicity in *Cdk12^KO^*-sgp53 allografts occurred despite absence of the focal tandem duplications implicated in neo-antigen formation in human prostate cancers with *CDK12* inactivation^18^ **(Fig S5C-D)**. Notably, however, luminal cells of the *Cdk12*-null prostate did display upregulation of numerous immunogenic pathways—including TNFA_signaling_via_NFkB, inflammatory response, IL6_JAK_STAT3_signaling, IL2_STAT5_signaling, and allograft rejection—suggesting other potential mechanisms underlying the observed lymphocytic infiltrate **(Fig S4D-E)**. In summary, *Cdk12* loss induces proinflammatory cytokine expression in prostate epithelial cells that corresponds with a T cell predominant immune infiltrate like that seen in human tumors^18^. This feature is recapitulated in our novel syngeneic *Cdk12^KO^*-sgp53 allograft model and corresponds with heightened sensitivity to immune checkpoint inhibitor therapy.

### Cdk12 loss mitigates progression of tumors with Pten inactivation

In contrast to the observed enrichment of *Trp53* sgRNA, our CRISPR screen revealed depletion of sgRNA targeting *Pten*, a gene rarely mutated in *CDK12*-mutant mCRPC **(Fig 3B)**. Given the mutual exclusivity of *CDK12* and *PTEN* inactivation in human tumors, we hypothesized that loss of *Cdk12*, an activator of mTOR signaling^35^, would mitigate progression of tumors driven by *Pten* inactivation. To test the hypothesis, we intercrossed our *Cdk12^pc-/-^* mice with animals from the well-known *Pten^f/f^* line—achieving simultaneous prostate-specific ablation of *Cdk12* and *Pten*. Strikingly, double knockout (*Pten^pc-/-^Cdk12^pc-/-^*) mice survived for a significantly longer time than mice with prostate-specific *Pten* ablation alone (*Pten^pc-/-^* mice) **(Fig 5A)**. Genitourinary tract weight was also significantly greater in *Pten^pc-/-^* mice versus *Pten^pc-/-^Cdk12^pc-/-^* mice at 52 weeks of age **(Fig 5B)**, corresponding with more aggressive tumors on gross observation **(Fig 5C)**. Histologically, prostates of *Pten^pc-/-^* mice demonstrated aggressive adenocarcinoma typical for the model, whereas those of *Pten^pc-/-^Cdk12^pc-/-^*mice showed maintenance of normal ductal morphology and markedly reduced stromal infiltrate **(Fig 5D)**. Similar findings were apparent in younger animals, as weights of individual prostate lobes were lower in *Pten^pc-/-^Cdk12^pc-/-^* versus *Pten^pc-/-^* mice at 24 weeks of age **(Fig 5E)**.

**Figure 5.**
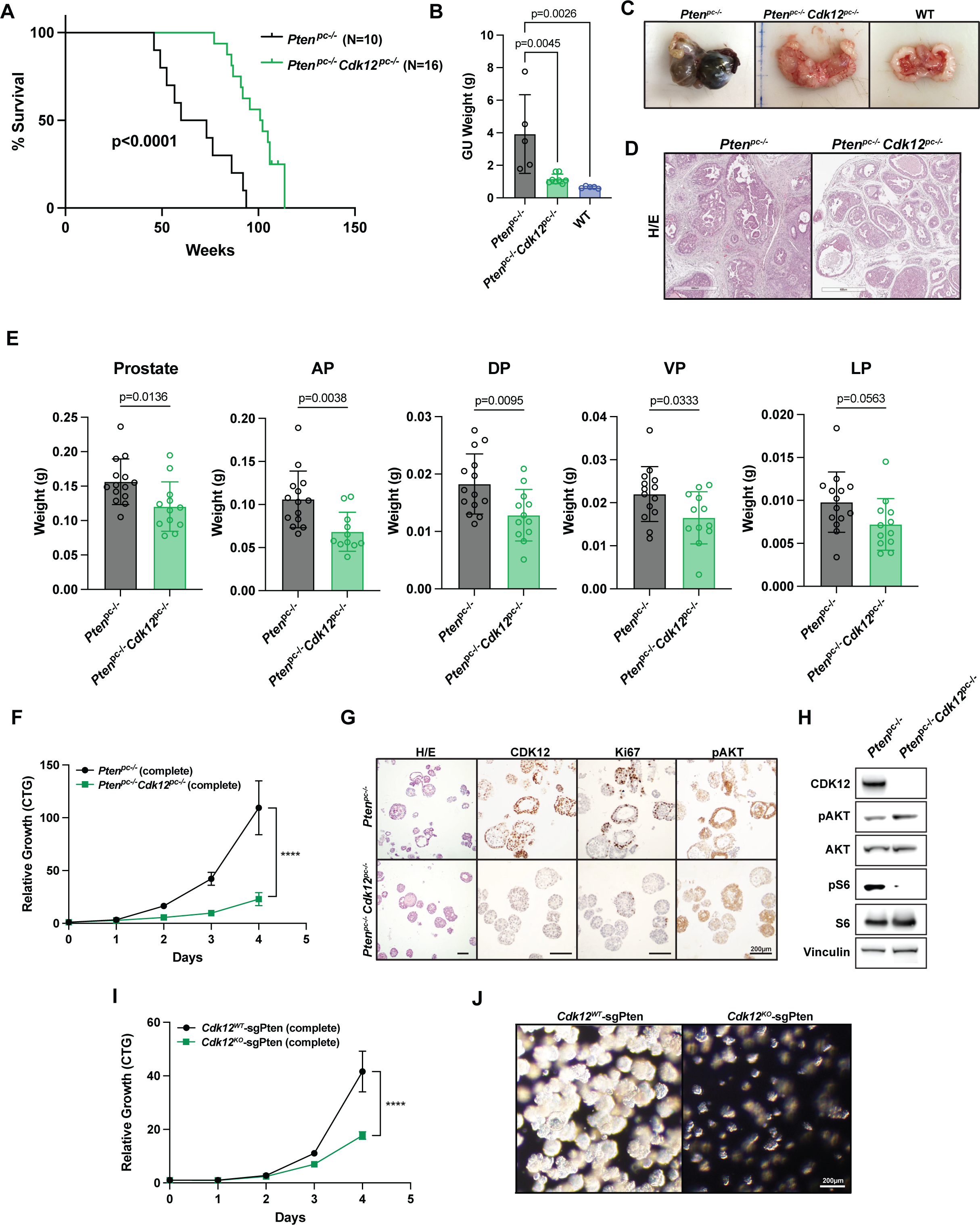
*Cdk12* ablation impairs tumor progression in the *Pten*-null mouse model of prostate cancer. **(A)** Kaplan-Meier plots demonstrating survival of prostate-specific Pten-null mice (*Pten^pc-/-^*) and *Pten^pc-/-^* with concomitant prostate-specific *Cdk12* ablation (*Pten^pc-/-^ Cdk12^pc-/-^*). **(B)** Genitourinary (GU) tract weights of *Pten^pc-/-^* and *Pten^pc-/-^ Cdk12^pc-/-^* mice at 52 weeks as well as wild-type mice (52 weeks). **(C)** Representative images of GU tracts from mice indicated in (B). **(D)** H/E-stained sections of *Pten^pc-/-^* and *Pten^pc-/-^ Cdk12^pc-/-^* prostate. Scale bars indicate 600μm. **(E)** Weights of whole prostate and individual lobes of *Pten^pc-/-^* and *Pten^pc-/-^ Cdk12^pc-/-^* mice at 24 weeks. **(F)** Cell proliferation in complete media of epithelial cell organoids derived from *Pten^pc-/-^* and *Pten^pc-/-^ Cdk12^pc-/-^* mice measured by CTG assay. (n= 4 samples per group). **(G)** Immunohistochemical staining of CDK12, Ki67, and phosphorylated AKT (pAKT) in cross sections of organoids described in (F). Scale bars indicate 200μm. **(H)** Protein expression of CDK12, pAKT, and pS6 in *Pten^pc-/-^* and *Pten^pc-/-^ Cdk12^pc-/-^* organoids with vinculin serving as a loading control. **(I)** Cell proliferation of basal cell-derived *Cdk12^WT^* and *Cdk12^KO^* organoids subjected to CRISPR-mediated *Pten* ablation (sgPten) as measured by CTG assay. (n= 4 samples per group) **(J)** Phase contrast images of organoids described in (I). Data are represented as mean ± SEM. Log-rank (Mantel-Cox) test was used to detect significance in (A). One-way ANOVA test was used to detect significance in (B). Unpaired t test was used for tumor weight in (E). Two-way ANOVA test was used for (F) and (I) ****p<0.0001.

We next aimed to determine whether the protective effect of *Cdk12* ablation in prostate cancer driven by *Pten* loss could be recapitulated *in vitro*. To do so, we generated prostate epithelial organoids from 24-week-old *Pten^pc-/-^* and *Pten^pc-/-^Cdk12^pc-/-^* mice, observing growth of the latter to be significantly blunted **(Fig 5F)**. Histologically, *Pten^pc-/-^Cdk12^pc-/-^* organoids displayed the absent-lumen phenotype but also demonstrated reduced cell proliferation (Ki67 staining) in the setting of equivalent Akt phosphorylation **(Fig 5G)**. Consistent with the established positive regulation of mTOR signaling by CDK12^35^, phosphorylation of S6 was markedly reduced in *Pten^pc-/-^Cdk12^pc-/-^* versus *Pten^pc-/-^* organoids **(Fig 5H)**. We confirmed these findings by using CRISPR-Cas9 to ablate *Pten* in the basal cell-derived *Cdk12^WT^*and *Cdk12^KO^* organoid lines described above. In agreement with the aforementioned findings, *Cdk12^KO^*-sgPten organoids demonstrated reduced growth versus *Cdk12^WT^*-sgPten organoids **(Fig 5I-J)**. These data reveal that *Cdk12* ablation impairs the growth of prostate cancer driven by *Pten* loss. This is consistent with findings from human samples, which demonstrate near mutual exclusivity between inactivating *CDK12* and *PTEN* mutations in mCRPC^36^ and emphasizes the context-dependent nature of cancer progression.

### CDK12 loss renders prostate epithelial tumor cells sensitive to CDK13 paralog inhibition

Our findings demonstrate that prostate cancer driven by *Cdk12* inactivation can effectively be modeled in mice—in both *in vivo* and *in vitro* systems. As in human mCRPC, *CDK12* inactivation is coupled with a T cell predominant immune infiltration and dependent on background mutational context.

To study the effects of CDK12 dysfunction in human cells, and to identify candidate synthetic lethal effects associated with CDK12 dysfunction, we used a previously described CRISPR-Cas9 engineered HeLa cell line, CDK12^as^ cells (“analogue sensitive”), where the only functional allele of the *CDK12* gene contains a kinase domain missense mutation rendering it sensitive to inhibition by a cell-permeable adenine analog, 1-NM-PP1 (1NM)^37^ **(Fig S6A)**. We first validated CDK12^as^ cells, demonstrating that 1NM administration reduced phosphorylation of serine 2 in the RNA polymerase II C-terminal domain (Pol-II CTD), reduced the ability to mount a nuclear RAD51 foci response to ionizing radiation, and caused PARP inhibitor sensitivity (**Fig S6B-G**). As expected, re-expression of wild-type CDK12 in CDK12^as^ cells reversed PARPi sensitivity (**Fig S6H-I**). We reasoned that other genes dysregulated in *CDK12* mutant cancers might be CDK12 synthetic lethal genes, and, therefore, screened the CDK12^as^ cells with a siRNA library targeting 297 candidates genes (**Table S1**, **Fig S6J-L** and see Methods) including genes involved in either mRNA splicing and/or the control of intronic truncating mutations, two processes where CDK12 dysfunction has been implicated^13^, genes whose expression is dysregulated in *CDK12* mutant compared to *CDK12* wild-type ovarian or prostate cancers^4,18,19^, genes that encode proteins with some evidence of interaction with CDK12^38^, and genes that encode proteins previously implicated as phosphorylation targets of CDK12^39^. Strikingly, this screen identified the siRNA encoding *CDK13* as the most deleterious for survival of 1NM-treated CDK12^as^ cells **(Fig 6A)**. We confirmed the ability of multiple *CDK13* siRNAs to induce cell death when CDK12 was inhibited by 1NM **(Fig 6B-C)**. CDK13 is a closely related paralog of CDK12, and the two proteins fulfill redundant functions in promoting Ser2 phosphorylation in the Pol II CTD.

**Figure 6.**
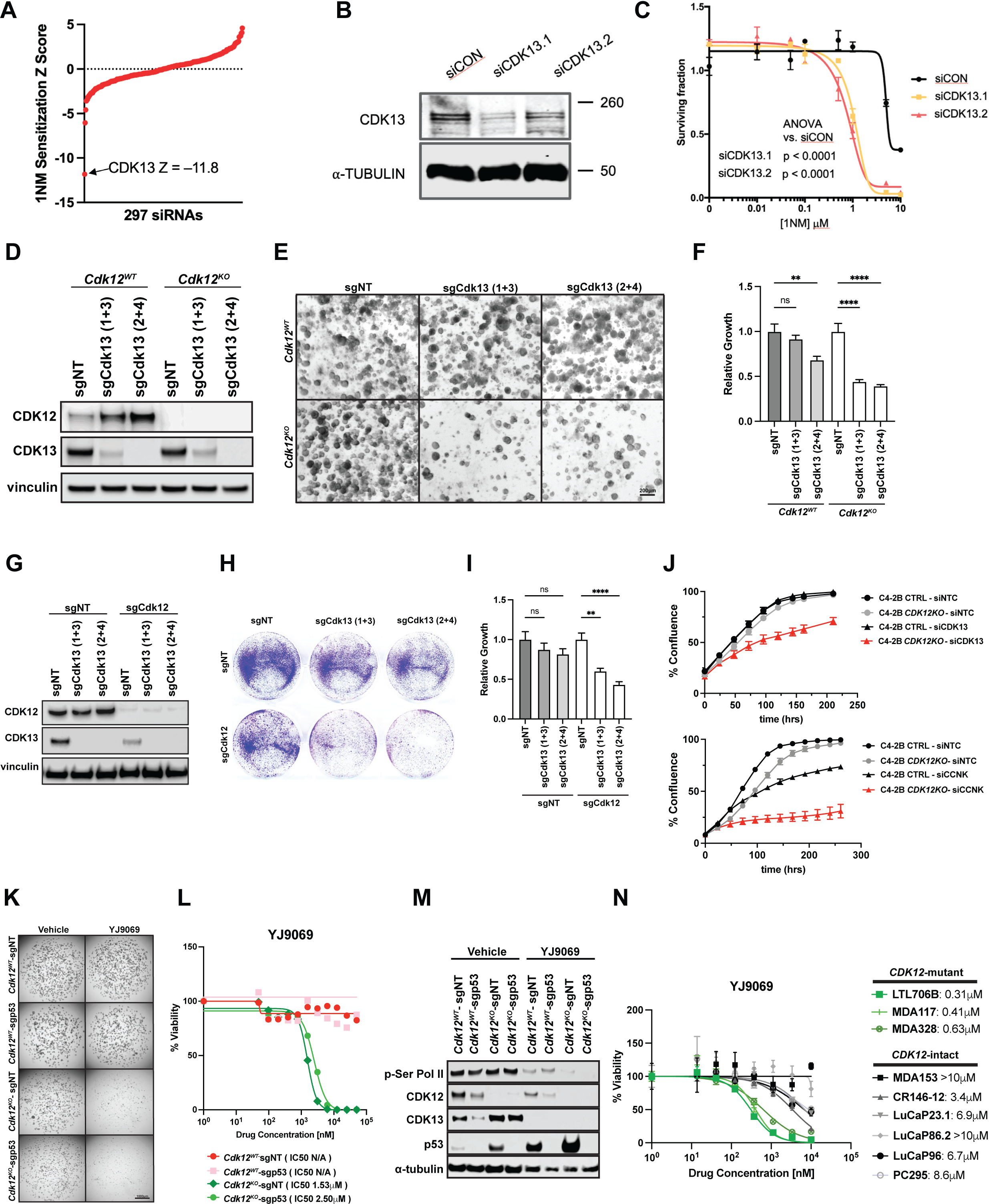
*Cdk12^KO^* organoids and *CDK12*-mutant tumors are preferentially sensitive to a CDK13/12 degrader. **(A)** Snake plot representing data from a siRNA screen to identify CDK12 synthetic lethal effects via 1NM sensitivity in CDK12^as^ cells. Negative Z scores indicate CDK12 synthetic lethal effects, with CDK13 representing the most profound effect. **(B)** Western blot indicating CDK13 gene silencing with two different siRNAs (siCDK13.1 and siCDK13.2. **(C)** Survival curve depicting cell survival in 1NM-exposed CDK12^as^ cells transfected with one of two unique CDK13 siRNAs (siCDK13.1 and siCDK13.2) or control siRNA (siCON). **(D)** CRISPR-mediated *Cdk13* (sgCdk13(1+3), or sgCdk13(2+4)) knockout in *Cdk12^WT^* and *Cdk12^KO^* organoids harvested on Day 5 after lentiviral transduction. Protein expression of CDK12 and CDK13 in organoids with vinculin serving as a loading control. **(E)** Brightfield images of organoids described in (D). Scale bars indicate 200μm. **(F)** Quantification of relative growth from images in (D). (n= 3 samples per group). **(G)** CRISPR ablation of *Cdk12* (sgCdk12) and *Cdk13* (sgCdk13(1+3), or sgCdk13(2+4)) in Myc-CaP cells. Protein expression of CDK12 and CDK13 in Myc-CaP cells treated with guide RNAs. **(H)** Colony formation assay showing cell survival in cells treated with indicated sgRNAs or control sgRNA (sgNT). (Representative data from 3 unique experiments) **(I)** Quantification of relative growth from images in (H). (Analysis of 11 high powered fields per sample over 2 unique experiments). **(J)** (Top panel) C4-2B prostate cancer cells subjected to CRISPR-based *CDK12* ablation (*CDK12*KO) or control sgRNA (C4-2B CTRL): Percent confluence measurement in the setting of siRNA-based *CDK13* knockdown (si*CDK13*) or treatment with control siRNA (siNTC). (Bottom panel) C4-2B *CDK12*KO and C4-2B CTRL cells: Percent confluence measurement in the setting of siRNA-based knockdown of *CCNK* (encoding cyclin K), or control siRNA treatment. (n= 3 samples per group). **(K)** Images of *Cdk12^WT^* and *Cdk12^KO^* with or without *Trp53* ablation organoids following treatment with CDK12/13 degrader (YJ9069). sgp53 indicates *Trp53* ablation, while sgNT indicates intact *Trp53*. Scale bars indicate 1000μm. **(L)** Viability curves and IC50 values of YJ9069 for groups described in (K). (n= 4 samples per group) **(M)** Protein expression of p-Ser RNA Pol-II, CDK12, CDK13, and p53 in *Cdk12^WT^* and *Cdk12^KO^* organoids with or without *Trp53* ablation subjected to YJ9069 degrader or vehicle treatment. **(N)** IC50 of organoids derived from patient-derived xenograft (PDX) lines with WT *CDK12* (MDA153, MDA146-12, LuCaP23.1, LuCaP86.2, LuCaP96, PC295) and inactivating *CDK12* mutation (LTL706B, MDA117, MDA328). (n= 3 replicates per line). Data are represented as mean ± SEM. One way ANOVA for multiple comparisons, two way ANOVA for multiple variables, **p<0.01, ****p<0.0001.

We then asked whether our *Cdk12^KO^* organoids were also susceptible to this paralog-based synthetic lethality. Indeed, CRISPR-based ablation of *Cdk13* in *Cdk12^WT^* and *Cdk12^KO^*organoids preferentially impaired organoid growth in the absence of *Cdk12* **(Fig 6D-F)**. We next developed an isogenic model of *Cdk12* loss by employing CRISPR-Cas9 to ablate *Cdk12* in Myc-CaP cells **(Fig 6G).** While *Cdk12* loss reduced cell number on colony formation assay, cell survival was further inhibited with co-ablation of *Cdk12* and *Cdk13* **(Fig 6H-I)**. We also employed CRISPR to ablate *CDK12* in the C4-2B prostate cancer cell line. When subjected to siRNA-mediated *CDK13* knockdown, these *CDK12* knockout cells exhibited more substantial growth reduction than si*CDK13*-treated C4-2B cells with intact *CDK12* **(Fig 6J)**. Knockdown of the transcript encoding cyclin K (*CCNK*)—the obligate binding partner required for kinase activity of both CDK12 and CDK13—also preferentially inhibited proliferation of *CDK12* knockout C4-2B cells **(Fig 6J)**. Taken together, these data substantiate CDK12 and 13 as synthetic lethal paralogs^28^.

While selective CDK13 inhibitors/degraders are in development, we employed a novel CDK13/12 degrader, YJ9069, in our organoid systems with the idea that active CDK12/13 dosage would be higher in wild-type cells versus *Cdk12^KO^* cells –thus providing a therapeutic window for preclinical studies and exploring paralog redundancy^40^. Treatment with YJ9069 markedly reduced the viability of *Cdk12^KO^*-sgNT and *Cdk12^KO^*-sgp53 organoids compared to *Cdk12^WT^*-sgNT and *Cdk12^WT^*-sgp53 organoids **(Fig 6K-L)**. Protein expression analysis showed reduction in Ser Pol II phosphorylation in YJ9069-treated organoids with intact *Cdk12*. Strikingly, phosphorylated Pol II was essentially absent in YJ9069-treated *Cdk12^KO^* organoids **(Fig 6M)**. Similar to the YJ9069 degrader, CDK13/12 inhibitors (YJ5118 and THZ531) exhibited more potent effects on cell viability in *Cdk12^KO^* organoids compared to *Cdk12^WT^* **(Fig S7A)**. In addition, *CDK12* knockout C4-2B cells showed preferential susceptibility to THZ531 treatment over C4-2B cells with intact *CDK12* **(Fig S7B-C)**. Finally, we analyzed the efficacy of CDK13/12 degrader therapy in our *Cdk12*-null Myc-CaP cells. In this system, YJ9069 effectively inhibited growth of sgCdk12 Myc-CaP cells (IC50 3.4 μM) but had no impact on Myc-CaP cells treated with control sgRNA (sgNT), even at a concentration of 20 μM **(Fig S7D)**. These results underscore the effectiveness of paralog-based lethality in cells lacking *Cdk12*.

Next, we analyzed the impact of YJ9069 treatment on three patient-derived xenograft (PDX) lines with biallelic inactivating mutations in *CDK12* (LTL706B, MDA117, and MDA328) **(Fig S8A)**. We successfully grew tumor chunks from LTL706B cells in the renal capsules of mice and confirmed positivity for prostate cancer markers (AR, KRT8, and PSMA) and absence of CDK12 by immunohistochemistry **(Fig S8B)**. We then propagated LTL706B tumors for six months, isolated tumor cells, and generated an organoid line for *in vitro* analysis **(Fig S8C)**. MDA117 and MDA328 are established PDXs obtained from the MD Anderson Prostate Cancer Patient-derived Xenograft Series (MDA PCa PDX) from which we developed organoid models. Treatment of each *CDK12*-mutant line with YJ9069 revealed IC50 values (0.31-0.63 μM) to be lower by an order of magnitude versus those of several established PDX lines with intact *CDK12* (MDA153, MDA146-12, LuCaP23.1, LuCaP86.2, LuCap96, PC295) **(Fig 6N, Fig S8A)**.

We then sought to determine whether CDK13 degradation was also effective in mice harboring *Cdk12*/*Trp53* loss. Indeed, intravenous administration of YJ9069 significantly blunted growth of subcutaneous *Cdk12^KO^*-sgp53 allografts but not allografts from the established TRAMP-C2 model **(Fig 7A-B)**. Similarly, sgCdk12 Myc-CaP allografts demonstrated significantly decreased tumor growth and tumor weights when treated with YJ9069; conversely, sgNT Myc-CaP allografts were insensitive to YJ9069 **(Fig 7C-D)**. In this model, tumor size reduction was coupled with increased frequency of apoptotic cells (TUNEL staining) in sgCdk12 Myc-CaP allografts treated with YJ9069, whereas no change in apoptosis was observed in YJ9069-treated sgNT Myc-CaP allografts **(Fig 7E-H)**. Treatment of mice harboring the same Myc-CaP allografts with the orally bioavailable CDK13/12 degrader YJ1206 phenocopied these data **(Fig S8D-E)**.

**Figure 7.**
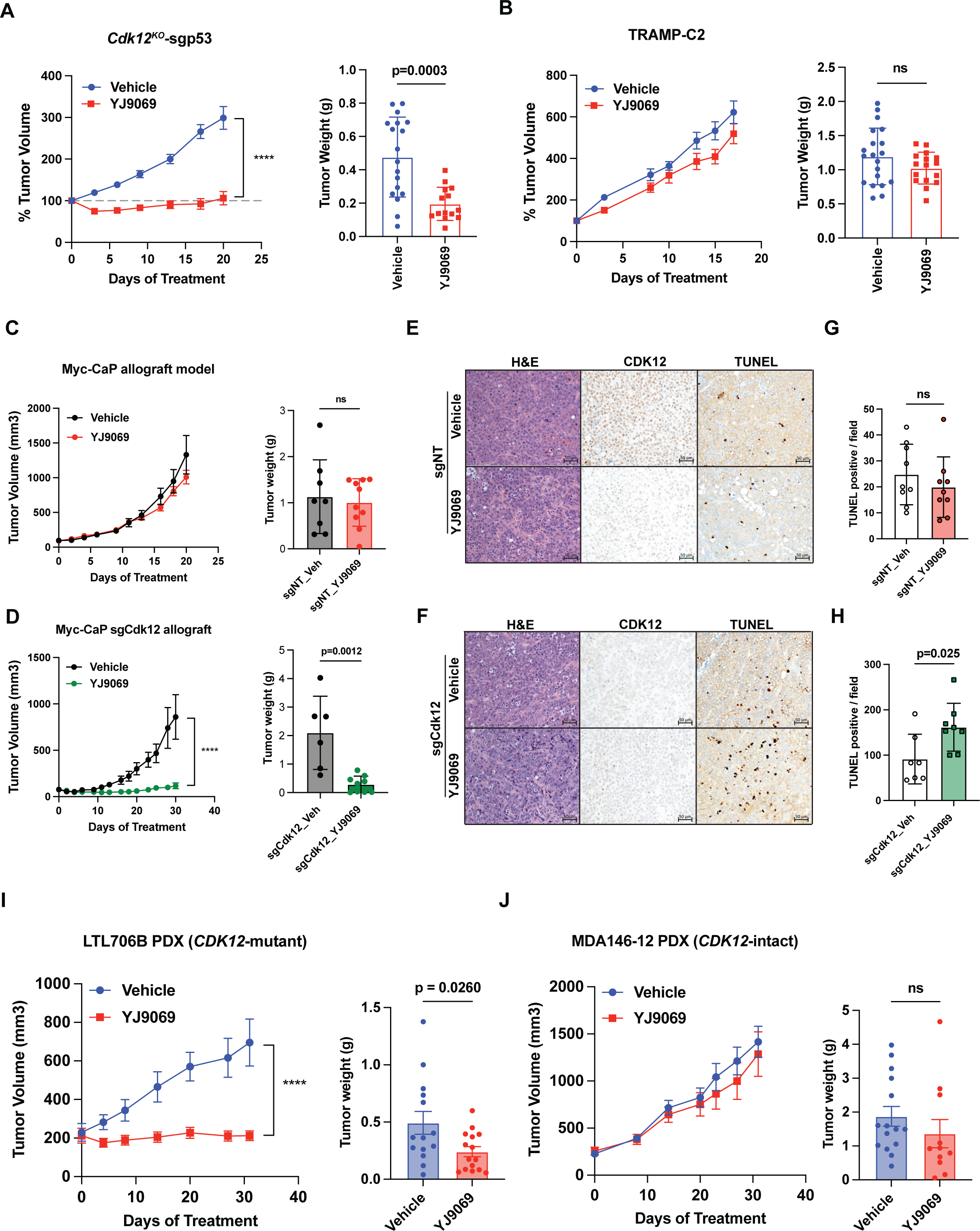
CDK13/12 degrader inhibits *CDK12*-mutant tumor growth *in vivo*. **(A)** *In vivo* treatment of *Cdk12^KO^*-sgp53 allografts with YJ9069 or vehicle: line graph indicates tumor volume normalized to baseline. Bar graph indicates tumor weight at endpoint. (n=9-10 mice, each with 2 tumors, per group) **(B)** *In vivo* treatment of TRAMP-C2 allografts with YJ9069 or vehicle: graphs as indicated in (A). (n=9-10 mice, each with 2 tumors, per group) **(C-D)** Unmodified (sgNT-treated) Myc-CaP allografts (C) or sgCdk12-treated Myc-CaP allografts (D) subjected to *in vivo* YJ9069 treatment: Line graphs indicate tumor volume. Bar graphs indicate tumor weight at end of treatment time course. (n=6-9 mice per group) **(E-H)** CDK12 immunohistochemistry and TUNEL staining of unmodified (sgNT-treated) Myc-CaP allografts (E) and sgCdk12-treated Myc-CaP allografts (F). Bar graphs (G-H) indicate quantification of TUNEL(+) cells per high powered field. Scale bars indicate 50μm. **(I-J)** YJ9069 treatment of subcutaneously implanted PDX lines, LTL706B (*CDK12*-mutant) and MDA146-12 (intact *CDK12*). Graphs indicate tumor volume over the treatment time course. (n=8-9 mice, each with 2 tumors per group) Two-way ANOVA test was used to detect significance in tumor volume of (A), (D), and (I), and unpaired t test was used for tumor weight in (A)-(D) and (I)-(J) and TUNEL staining (G)-(H). ****p<0.0001; ns, not significant.

Finally, we aimed to determine if *CDK12*-mutant PDX tumors shared the same *in vivo* sensitivity to CDK13/12 degrader therapy. Indeed, we observed that subcutaneous allografts derived from *CDK12*-mutant LTL706B were highly sensitive to YJ9069 therapy (**Fig 7I**). On the contrary, those derived from *CDK12*-intact MDA146-12 showed similar growth in the setting of YJ9069 and vehicle treatment **(Fig 7J)**. Thus, human *CDK12*-mutant prostate cancer PDXs are preferentially sensitive to a CDK13/12 degrader as compared to *CDK12* wild-type PDXs. Together, these data demonstrate that *CDK12* loss increases dependence on CDK13 to render prostate cancer cells sensitive to CDK13/12 inhibitors and degraders, such as YJ9069. These findings suggest that this vulnerability may be leveraged clinically in the future in *CDK12*-mutant cancers, ideally with CDK13-selective inhibitors/degraders; however, our data suggest that a therapeutic window can be achieved even with dual CDK13/12 degraders/inhibitors.

## DISCUSSION

Through multiple *in vivo* models, this study demonstrates that *Cdk12* is a tumor suppressor gene, the inactivation of which is sufficient to induce DNA damage and immunogenic preneoplastic lesions in the mouse prostate. Since *Cdk12* is required for both genomic stability and cell proliferation, the impact of its loss is contingent on background mutational status. With concomitant knockout of the DNA damage response gene *Trp53*, *Cdk12* ablation promotes tumorigenesis. Conversely, because CDK12 promotes cell growth and proliferation—in part through activation of mTOR signaling^35^—its absence inhibits growth of *Pten*-null prostate tumors. Finally, CDK12 and its paralog, CDK13, play redundant roles to facilitate cell proliferation and survival. This genetic interaction renders *CDK12*-null tumors uniquely sensitive to CDK13 degradation—a form of paralog-based synthetic lethality that can be applied in future clinical pharmacotherapy.

Clinical data amassed over the last decade suggests that *CDK12* functions as a tumor suppressor gene in multiple adenocarcinomas including serous ovarian carcinoma^20,41,42^ and triple-negative breast cancer^43^. In prostate cancer, our group identified a putative tumor suppressor role for *CDK12* through whole-exome sequencing of samples compiled from SU2C^21^, Mi-Oncoseq^22^, and University of Michigan rapid autopsy cases^23^. While these correlative findings established a relationship between *CDK12* inactivation and aggressive prostate cancer, it remained unclear whether *CDK12* loss *per se* could induce prostate tumorigenesis or progression. We addressed this question here, finding that conditional ablation of *Cdk12* (with specific targeting of the kinase domain) in the mouse prostate epithelium is sufficient to induce preneoplastic lesions. These data reveal a *bona fide* tumor suppressor role for *Cdk12* in prostate cancer. In our mouse model, prostate lesions assume histological subtypes including focal HGPIN and AIP, while exhibiting increased basal cell content. Organoids derived from the *Cdk12^KO^* prostate recapitulated the phenotype, demonstrating an abnormal compact morphology seen in the setting of other canonical mCRPC mutations^31^. Strikingly, *Cdk12^KO^*organoids exhibited a basal-to-luminal differentiation defect (in agreement with the basal cell excess observed *in vivo*) as well as transcriptomic changes consistent with those seen in human *CDK12*-mutant mCPRC^18^. Consistent with the high prevalence of castration resistance among *CDK12*-mutant clinical samples^18^, *Cdk12^KO^* organoids displayed increased AR target gene expression, as well as resistance to androgen depletion and enzalutamide therapy. Taken together, our *in vivo* and *ex vivo* systems define *Cdk12* as a tumor suppressor gene while reliably modeling key aspects of human disease.

The tumor suppressor function of *CDK12* has been attributed to its regulation of DNA damage responses. For instance, in ovarian cancer, *CDK12* loss causes downregulation of transcripts in the homologous recombination DNA repair pathway—a phenotype attributed to reduced CDK12/cyclin K-mediated transcriptional elongation of the corresponding genes^44^. Aligned with these findings, *CDK12* has been defined as positive regulator of *BRCA* expression in both ovarian^45^ and triple negative breast cancer^46^. Conversely, Popova et al. defined a unique defect characterized by numerous focal tandem duplications and intact homologous recombination in the 4% of serous ovarian cancers lacking functional *CDK12*^20^. The same held true in our mCRPC exome sequencing study, as *CDK12*-mutant tumors constituted a unique mCRPC subtype, genetically distinct from those with other primary genetic drivers, including homologous recombination deficiency. *CDK12*-mutant tumors were characterized by a genomic instability in which recurrent gains secondary to focal tandem duplications yielded putative neo-antigens^18^.

In our mouse model, *Cdk12* loss was sufficient to induce DNA damage, as indicated by γH2AX immunohistochemistry. Conspicuously, γH2AX(+) lesions geographically corresponded with p53 protein expression not observed elsewhere in the tissue, suggesting *Cdk12*-null cells to have increased dependence on p53 for maintenance of genomic stability. *Cdk12^KO^*organoids similarly exhibited elevated γH2AX, and concomitant increases in p53 and its target genes. Strikingly, unbiased *in vivo* CRISPR screening identified *Trp53* as the most enriched sgRNA in allograft tumors derived from *Cdk12^KO^* organoids. In line with our *in vivo* and organoid data, this finding indicated that tumorigenesis in the setting of *Cdk12* loss could be enhanced with concomitant loss of a compensatory DNA damage response gene. Indeed, while *Cdk12^KO^* organoid cells did not form viable allografts when injected subcutaneously into mice, those with bigenic *Cdk12/Trp53* loss formed tumors. These data broadly agree with clinical sequencing data, in which *CDK12* and *TP53* inactivation often occur in the same tumors^18^.

Biallelic *CDK12* loss in human mCRPC engenders a T cell-predominant immune response that may in part be driven by neo-antigens arising from translated products of focal tandem duplications^18^. In our *Cdk12^KO^* mouse model, we observed preneoplastic prostate lesions to be similarly infiltrated by CD4(+) and CD8(+) T cells to a considerably greater degree than F4/80(+) macrophages or NK1.1(+) natural killer cells. Strikingly, a nearly identical infiltrate was phenocopied upon subcutaneous transplantation of *Cdk12*/*Trp53* bigenic mutant organoid cells into immunocompetent hosts. The composition of the immune infiltrate distinguished our newly generated syngeneic model from preexisting Myc-CaP and TRAMP-C2 systems with wild-type *Cdk12*. Both of these allograft types induced few CD4(+) and no CD8(+) cells. Indeed, we are unaware of other syngeneic models capable of promoting significant CD8 infiltration. As such, *Cdk12*/*Trp53*-null allografts may in the future be broadly used to study T cell-driven tumor immunity. Moreover, the sensitivity of these tumors to immune checkpoint inhibitor therapy opens a promising clinical immunotherapy avenue.

Despite inducing a T cell-predominant immune response like that seen in human tumors, *Cdk12*/*Trp53*-null allografts did not contain focal tandem duplications like those observed in human mCRPC and serous ovarian cancer^18,20^. Instead, we surmise the response in this system is driven by one or more of the numerous inflammatory pathways upregulated in *Cdk12^KO^* organoids. It is notable that the most upregulated HALLMARK signaling pathways in this model included those corresponding with TNFα, inflammatory, IL-6, complement, and IL-2 signaling. Further exploration of the impact of *Cdk12* loss on pro-inflammatory signaling is an important future direction that will be facilitated by our syngeneic model.

Despite its status as a tumor suppressor gene, *CDK12* has also been found to promote cell proliferation. For instance, using an analog-sensitive *CDK12* knock-in cell line enabling its rapid and specific pharmacologic inhibition, Chirackal and colleagues demonstrated that CDK12 promotes G1/S transition through enhancing RNA Pol-II processivity at key DNA replication genes^47^. Similarly, conditional ablation of *Cdk12* in mouse neural progenitors impairs their transit through the cell cycle^12^. Conversely, elevated CDK12 expression has been demonstrated in human cancers. This is best characterized in HER2(+) breast cancers, in which both genetic amplification and increased protein expression of CDK12 have been observed^43,48^. In HER2(+) breast cancer cell lines, CDK12-dependent alternative splicing has also been linked to increased invasiveness and metastatic potential^49^. Furthermore, elevated CDK12 protein levels are seen in gastric cancer and correlate with invasive histology and reduced patient survival^50^. Choi and colleagues recently elucidated a complicated reciprocal interaction between CDK12 and 4E-BP1 that promotes the translation of several mTORC1-dependent mRNAs critical for MYC-driven transformation and mitosis^35^. This latter report suggests that proliferation of prostate cancer cells dependent on mTOR (and upstream PI3 kinase) signaling may, unlike *Trp53*-null cells, exhibit growth inhibition in the setting of *Cdk12* loss.

We directly tested this premise by co-ablating *Cdk12* in the prostate epithelium of *Pten*-null knockout mice. When compared with *Pten*-null animals with intact *Cdk12*, these double knockout mice demonstrated improved survival and markedly reduced prostate tumor size. We recapitulated the growth defect induced by *Cdk12* loss in an *in vitro* organoid model and demonstrated this to correspond with reduced S6 phosphorylation—indicative of impaired mTOR signaling. These findings agree with our previously published whole-exome sequencing, which showed that *CDK12*/*PTEN* bigenic mutations occur very rarely in human mCRPC^18^.

CDK12 and CDK13 are evolutionarily related, structurally similar kinases, both of which phosphorylate the Pol II CTD to promote transcriptional elongation of overlapping target gene sets^16^. Fan et al. recently highlighted the functional redundancy of these paralogs, applying CRISPR technology to generate analog sensitive CDK12 and CDK13 proteins in leukemia cell lines. This system demonstrated that, while CDK12 and CDK13 regulate transcription of some genes independently, only dual inhibition of both kinases is sufficient to induce genome-wide transcriptional changes and loss of Pol II CTD phosphorylation, as well as associated proliferation defects and cell death^17^. These findings are in line with data from ovarian cancer cell lines showing the therapeutic promise of the dual CDK12/13 inhibitor THZ531^51,52^.

Given the functional redundancy of CDK12 and CDK13 in maintaining cell proliferation and survival, we hypothesized that *Cdk12*-null cells would show enhanced sensitivity to CDK13/ 12 inhibitor/degrader therapy over cells with intact *Cdk12*. We first demonstrated this *in vitro*, determining that CRISPR-mediated *Cdk13* ablation has a greater impact on cell number in *Cdk12^KO^* versus *Cdk12^WT^* organoids. We next used YJ9069, a novel CDK13/12 degrader, to treat several mouse-derived cell and organoid lines lacking *Cdk12*. *In vitro*, YJ9069 IC50 values were significantly lower in *Cdk12*-null cell lines. The same was true *in vivo*, as *Cdk12* knockout allografts demonstrated comparatively reduced tumor growth and increased tumor cell apoptosis with YJ9069 treatment. Strikingly, the same premise held in PDXs, as human mCRPC lines with biallelic *CDK12* inactivation also showed enhanced sensitivity to CDK12/13 degradation. As such, YJ9069 and related molecules are agents with promising clinical applicability in *CDK12*-mutant prostate cancer.

Combined, findings from this study define the role of CDK12 in prostate cancer while generating valuable murine models of *Cdk12* loss that recapitulate human *CDK12*-mutant prostate cancer. *Cdk12* is identified as a tumor suppressor gene that works in a context-dependent manner to promote prostate cancer characterized by enhanced T cell infiltration. The amassed findings from this study have the potential for near-term clinical impact for patients with *CDK12/TP53*-mutant prostate cancers, whereby immune checkpoint blockade therapies are likely to elicit an enhanced response. Furthermore, CDK13/12 or CDK13-specific inhibitors have strong potential as possible treatment options for *CDK12*-mutant prostate cancer.

### Limitations of the study

While we aimed to comprehensively model prostate cancer characterized by *CDK12* loss, further study into its connection with AR signaling will be a fruitful area of future inquiry. Similarly, while we presented the mitigating effect of *Cdk12* ablation on tumor progression in the *Pten* knockout mouse model as a counterpoint to its effect on cancers driven by *Trp53* loss, a better mechanistic understanding of this phenomenon—and the degree to which it represents another form of synthetic lethality—will also be valuable to pursue. While *Cdk12* ablation in the mouse prostate induces gene expression alterations and T cell infiltration like that observed in human *CDK12*-mutant tumors, the focal tandem duplications seen in human tumors were, thus far, not seen in murine tumors in which *Cdk12* was ablated. Whether these tumors eventually develop focal tandem duplications with aging will be of tremendous interest, as well as defining the mechanism as to how these focal tandem duplications are formed. Finally, we posit that reduced CDK13 action underlies the uniquely antagonistic effect of CDK13/12 inhibitors and degraders on *CDK12-*mutant prostate cancer. While no CDK13-specific inhibitor/ degrader currently exists, we surmise that such an agent will have great utility for patients with mCRPC characterized by *CDK12* inactivation by enhancing the therapeutic window as well as fully taking advantage of paralog-based synthetic lethality.

## ACKNOWLEDGEMENTS

We acknowledge Shuqin Li, Derrick Ekanayake, Fengyun Su, and Rui Wang for their technical assistance and Brian Magnuson for his support on sequence analysis from the CRISPR library screening experiment. We also thank Lisa McMurry, Amanda Miller, and Christine Caldwell-Smith for their histology support. We thank Dr. Arno Greenleaf at Duke University for provision of the CDK12^as^ cell line. We dedicate this study to the memory of Dr. Nora Navone for her development of the MD Anderson PDX library, from which we received several samples for this study. Her legacy continues with the efforts of her trainee, Dr. Estefania Labanca who is a co-author on this study. This work was funded by the Prostate Cancer Foundation (PCF), a National Cancer Institute (NCI) Prostate Specialized Programs of Research Excellence (SPORE) Grant (P50-CA186786), the NCI Early Detection Research Network (EDRN) (U2C-CA271854), an NCI Outstanding Investigator Award (R35-CA231996, A.M.C.), and a Programme Grant from Cancer Research UK to C.J.L. (DRCRPGNov21y100001). J.C.T. is supported by a Department of Defense Prostate Cancer Research Program Idea Development Award (W81XWH-21-1-0458). A.M.C. is a Howard Hughes Medical Institute Investigator, A. Alfred Taubman Scholar, and American Cancer Society Professor.

## AUTHOR CONTRIBUTIONS

J.C.T., J.C., F.Y.F., C.J.L, K.D., and A.M.C. conceived the study and designed the experiments. J.C.T., Y. Cheng, and P.S. performed all *in vitro* and *in vivo* functional experiments with assistance from Y. Chang, S.E., A.P., L.X., and A.J.T. Y.W., D.R.R., and X.C. generated sequencing libraries and performed the sequencing. Xiao-Ming W. performed *in situ* hybridization and immunofluorescence staining. R.M. carried out histopathological evaluations. Y.Z., C.C., R.J.R., and M.C. performed bioinformatic analyses. Y.B. conducted immune profiling by flow cytometry. J.N., R.B., and S.J.P. performed and analysed experiments involving CDK12^as^ cells, including siRNA screens and validation. C.J.L. supervised experiments involving CDK12^as^ cells. J.C. generated C4-2B *CDK12 KO* line and performed experiments under supervision of F.Y.F. Y. Chang, J.Y., L.Z., Z.W., Xiaoju W., and K.D. were involved in the discovery, synthesis, and initial profiling of YJ9069, YJ1206, and YJ5118 compounds. Y.W. and E.L. provided PDX lines. J.C.T., C.J.L., and A.M.C. wrote the manuscript and organized the final figures. S.J.M. reviewed and edited the manuscript.

## METHODS

### Mice

*Cdk12^f/f^* mice (B6.129-*Cdk12 ^tm1Fmj^*/Narl) were obtained from the National Laboratory Animal Center (Taiwan, R.O.C.). *Probasin-Cre* (Stock#026662), *Rosa^mT/mG^*(Stock#007676), and *Pten^f/f^* (Stock#006440) mice were obtained from the Jackson Laboratory (Bar Harbor, ME). The animals were interbred, backcrossed, and maintained on a C57Bl/6J background. For syngeneic models, six-to-eight-week-old male C57Bl/6J (Stock# 000664) mice were obtained from the Jackson Laboratory, and male FVB (Stock #207) mice were obtained from Charles River Laboratory (Wilmington, MA). NOD Cg-Prkdc<SCID> ll2rg<TM1WJL>SzJ (NSG) mice were obtained from the Jackson Laboratory, (Stock#005557) while CB17SCID mice were obtained from Charles River (Stock#236). All animals were housed in pathogen-free containment with a 12-hour light-dark cycle and *ad libitum* food and water. The University of Michigan Institutional Animal Care and Use Committee (IACUC) approved all animal studies.

### Histological analysis and immunohistochemistry

Prostate tissue and allograft tumors were fixed in formalin overnight, dehydrated with ethanol, and paraffin embedded. Five μm-thick sections were prepared for H&E staining and immunohistochemistry. Two pathologists with expertise in genitourinary evaluated the H&E-stained formalin-fixed paraffin-embedded (FFPE) tissue sections in a blinded manner. Before the assessment, four histopathological scoring schemas were created based on the temporal progression of prostate pathology. These categories were: Category 0: Normal prostatic epithelium; Category 1: Epithelial hyperplasia; Category 2: Focal high-grade prostatic intraepithelial neoplasia (PIN); Category 3: Florid high-grade PIN/atypical intraepithelial neoplasm (AIP) and intraductal carcinoma. Each prostate sample was evaluated for the overall percentage prevalence for each of these four categories. Immunohistochemistry was performed manually or using the Ventana automated slide staining system (Roche-Ventana Medical System). Antibody concentrations are listed in **Table S2**. For immunohistochemical staining of organoids, organoids were embedded in Histogel and fixed with 4% PFA for 1h, then ethanol dehydrated, and paraffin embedded. For manual staining procedure, samples were then deparaffinized and incubated in Antigen Unmasking Solution (Vector Laboratories, H-3300). Endogenous peroxidases were inactivated via incubation in 3% hydrogen peroxide (Sigma). Primary antibodies were diluted in 10% normal goat serum with overnight incubation; and antibody detection was achieved with species-specific VECTSTAIN Elite ABC kits (Vector Labs) and DAB Peroxidase Substrate kit (Vector Labs).

### RNA *in situ* hybridization

*Cdk12* gene expression was detected in FFPE tissue sections using the RNAscope 2.5 HD Brown kit (Advanced Cell Diagnostics, Newark, CA) and the target probe against the mouse *Cdk12* gene (cat # 444881). The *Cdk12* target probe is complementary to NM_02695.2, 102-1021nt. RNA quality was evaluated by a positive control probe against mouse low-copy housekeeping gene (*ppib*). Assay background was evaluated by a negative control probe targeting bacterial *DapB* gene. FFPE tissue blocks were cut into 4μm sections. The tissue sections were baked at 60°C for one hour, deparaffinized in xylene, and dehydrated in 100% ethanol followed by air drying. After hydrogen peroxide pretreatment and target retrieval in citrate buffer at 100°C, tissue sections were permeabilized using protease and hybridized with the target probe in the HybEZ oven for 2 hours at 40°C, followed by a series of signal amplification steps. Finally, the sections were chromogenically stained with DAB and counterstained with 50% Gill’s Hematoxylin I (Fisher Scientific, Rochester, NY).

### Immunofluorescence

Immunofluorescence (IF) was performed on whole FFPE 5-µm-thick tissue sections using anti-TP63 mouse monoclonal antibody (1:100; Abcam, catalog no. ab735), anti-CK8 rabbit polyclonal antibody (1:100, Abcam, catalog no. ab53280), anti-p53 rabbit polyclonal antibody (Leica, cat no. P53-CM5P-L), and anti-CDK12 rabbit polyclonal antibody (Atlas, cat no. HPA008038). TP63/CK8 double IF was carried out on a Discovery Ultra automated slide staining system (Roche-Ventana Medical Systems) using CC1 95°C for antigen retrieval, followed by first antibody (anti-TP63) incubation, OmniMap anti-mouse horseradish peroxidase (HRP), and signal development using Discovery Cy5 Kit (RTU, Roche-Ventana Medical Systems, catalog no. 760-238). The second antibody staining was performed consecutively with heat denaturation before the second primary antibody (anti-CK8) incubation, OmniMap anti-rabbit HRP kits, and signal development using Discovery FITC (RTU, Roche-Ventana Medical Systems, catalog no. 760-232). With a similar algorithmic process, p53/CDK12 double IF was performed first with p53 IF with Cy5, followed by CDK12 IF with FITC. The staining was independently assessed by three study participants including one pathologist (J. Tien, Xiao-Ming Wang, and R. Mannan) at ×100 and ×200 magnification to assess for presence and pattern of expression.

### Prostate isolation and flow cytometry

Mouse prostates were harvested from 52-week-old mice, and single cell isolation was adapted from previously published protocols^31,53^. First, prostates were digested with collagenase Type II (Gibco) for 1 hour at 37°C followed by TryLE (Gibco) digestion. After TryLE digestion, samples were inactivated with an excess of DMEM containing 10% fetal bovine serum (FBS), and samples were sequentially passed through 100 μm and 40 μm cell strainers to remove debris. Flow cytometry analysis used established marker profiles^31^. Briefly, fresh cells were incubated in PBS with fluorophore-conjugated antibodies at dilutions indicated in **Table S2** for 30 minutes at 4°C. DAPI was added for the final 5 minutes of the incubation to act as a dead cell marker. Cells were analyzed on MoFlo Astrios EQ running Summit software (version6.3; Beckman-Coulter, Brea, CA). Gates were established using fluorescence minus one approach, and plots were generated in FCS Express 7 (DeNovo Software). The Flow Cytometry Core from the University of Michigan assisted with the flow sorting experiment.

### Organoid culture/growth assay

Isolated prostatic epithelial cells were embedded in 50-μL drops of Matrigel and overlaid with mouse prostate organoid medium as previously described^53^. Media was changed every 2-3 days and organoids were passaged on a weekly basis. Prostate organoid cells were seeded at 1000 cell density in a matrix dome on Day 0 in medium without EGF or DHT. Cell viability was assayed starting at Day 1 for 6 days according to the CellTiter-Glo 3D kit (Promega G9683). For the antiandrogen response assay, the procedure was adapted from a previously published protocol^54^. Briefly, organoids were seeded at 2000 cells on Day 0 in media minus EGF, with 1 nM DHT or 10 μM of enzalutamide (Selleck Chemicals) added. Cell growth was assayed on Day 7 using the CellTiter-Glo 3D kit.

### Adenoviral Cre and CRISPR/Cas-9 lentiviral transduction

Adenoviral Cre-mediated recombination of *Cdk12* in mouse prostate organoids was performed by adenoviral delivery of CRE recombinase as previously described^55^. Similarly, CRISPR/Cas-9 mediated knockout of *Trp53*, *Pten*, and *Cdk13* was performed by lentiviral delivery of plasmids encoding Cas9 and gRNA sequences using LentiCRISPRv2 plasmids. sgRNA sequences are listed in **Table S2**.

### Allograft model

Organoid allograft models were generated by subcutaneous injection of Matrigel-suspended organoid cells (3 x 10^6^ cells per injection) into dorsal flanks of NSG mice. Animals were monitored for tumor growth weekly. Once *Cdk12^KO^*-sgp53 tumors reached 1000 mm^3^, the tumors were resected, cut into small chunks, and subcutaneously implanted into both flanks of C57Bl/6J mice for generation of the syngeneic model.

### Single cell RNA sequencing (scRNA-seq) and data analysis

scRNA-seq for dissociated mouse prostate tissues and organoids was performed using 10X Genomics Chromium Single Cell 3’ Library Gel bead Kit V3.1 according to the manufacturer’s protocol. The libraries were sequenced with the Illumina Hiseq 2500 or NovaSeq 6000 according to recommended specifications. After sequencing, read demultiplexing, alignment, and gene quantification were conducted with the 10X Genomics Cell Ranger pipeline (v5.0) and the pre-built mouse reference genome (mm10). For libraries from mouse prostate tissues, custom reference genome including sequences of GFP and tdTomato were used. Downstream analyses using the filtered gene count matrix were performed with R package Seurat (v4.1)^56^ if not specified otherwise. Low quality cells were further filtered based on total UMI, number of detected genes, and fraction of mitochondrial reads per cell using the Outlier function from the scatter package^57^; specifically, cells that were three times of mean absolute deviation (MAD) away from median on the three metrics were removed. In addition, putative doublets were identified with the R package scDblFinder^58^ and removed. After cell filtering, mitochondrial genes were also removed from the matrix. SoupX^59^ was used to adjust the count matrix in order to minimize impact of ambient RNA. After all QC steps, the SoupX corrected count matrix was then normalized using the NormalizeData function with the "LogNormalize" method. The top 2000 highly variable genes were then identified with FindVariableFeatures with the “vst” method, followed by ScaleData, RunPCA, and RunUMAP steps to obtain a 2-D map of the cells. The FindNeighbors and FindCluster functions were used to assign cells into clusters. Cell annotation was based on prediction using the TransferData method and a public dataset as reference^60^. RNA velocity analysis on the *Cdk12^WT^* organoid was conducted with velocyto^61^ to count spliced and unspliced RNA and scvelo^62^ to calculate RNA velocity and pseudotime and visualize velocity vector field as streamlines. Cells from *Cdk12^KO^* organoids were projected into UMAP of *Cdk12^WT^*and annotated using the MapQuery function of Seurat. To conduct GSEA between *Cdk12^KO^* and *Cdk12^WT^*, pseudo-bulk gene expression profiles were generated by summing counts for each cell type in *Cdk12^KO^* and *Cdk12^WT^*, respectively; normalized expression in TPM was then calculated with edgeR by incorporating TMM scaling factors^63^. Genes ranked by logFC were used as input for pre-ranked GSEA with fgsea^64^. Hallmark gene sets were downloaded from MsigDB^65^. The human CDK12 gene signature was defined using common genes up-regulated in prostate cancer patients with *CDK12* mutation and siCDK12 knockdown LNCaP cells^18^. The mouse homolog genes of the human CDK12 signature were mapped using biomaRt^66^.

### *In vivo* CRISPR screening

The MusCK library was a gift from Xiaole Shirley Liu (Addgene 174196). The MusCK library contains guide RNAs targeting 4922 mouse genes that are implicated in cancer. A total of 10^7 *Cdk12^KO^* organoids were transduced with lentivirus containing the MusCK library at a multiplicity of Infection (MOI) of 0.3 to achieve about 100x coverage. After puromycin selection for 5 days, ∼30% of the surviving cells were stored as Day0 input samples at -80°C, and the rest of cells were cultured for *in vivo* screening. 3 x 10^6 cells were prepared for each injection site for a total of 10 injection sites. Animals were monitored every week for tumor growth. Resulting tumors were harvested for genomic DNA extraction. PCR and purification of the regions containing the sgRNA were performed to generate the sequencing library. Each library was sequenced at approximately 3 million reads. Cutadapt^67^ was used to trim reads to the bare sgRNA sequences. The trimmed reads were then aligned to a reference built from the sgRNA sequences in the library using bowtie2(version 2.4.5)^68^. Finally, MAGeCK (version 0.5.9.5)^69^ was used to count the sgRNAs.

### RNA isolation and quantitative real-time PCR

Total RNA was isolated using QIAzol Lysis Reagent (QIAGEN), and cDNA was synthesized following Maxima First Strand cDNA Synthesis Kit (Thermo Fisher Scientific) instructions. Quantitative real-time PCR (qPCR) was performed in triplicate using either ThermoFisher Taqman Gene Expression assay or standard SYBR green protocols using SYBR™ Green PCR Master Mix (Applied Biosystems) on a QuantStudio 5 Real-Time PCR system (Applied Biosystems). The target mRNA expression was quantified using the ΔΔCt method and normalized to the expression of the housing keeping gene. Primer sequences and Taqman probes are listed in **Table S2.**

### PDX models

The LTL706B (*CDK12*-mutant) tumor was obtained from the Vancouver Prostate Centre and initially established in the renal capsule of NSG mice with a testosterone pellet (12.5 mg) implant. Once tumors grew successfully, we transferred them into subcutaneous pockets of CB17SCID mice for therapy studies. Other PDX, such as MDA117, 328 (*CDK12*-mutant) and MDA153, 146-12 (*CDK12*-intact), were obtained from MD Anderson. LuCaP23.1, 86.2, and 96 (*CDK12*-intact) PDX tumors were obtained from the University of Washington. Tumors from MDA and LuCaP PDX lines were maintained subcutaneously in dorsal flanks of CB17SCID male mice. The PC295 (*CDK12*-intact) PDX line was obtained from Erasmus Medical Center, Rotterdam, the Netherlands^70^.

### Compounds

YJ9069, YJ1206, and YJ5118 were synthesized in Dr. Ke Ding’s lab^71^. THZ531 was purchased from Cayman Chemical Company or Selleckchem. 1NM (aka 1NM-PP1) was purchased from Axon Medchem. Talazoparib was purchased from Selleckchem.

### Drug treatment of organoids and cell lines

To generate drug response curves, mouse organoids were digested with TryPLE for 10 min at 37°C, dissociated into single cells, and neutralized with FBS. Cells were resuspended in 20% Matrigel, plated in triplicate at a seeding density of 5000 cells/well in 48-well microplates. The next day, 8 doses of YJ9069 were dispensed at 3-fold dilution from 0.01 μM to 10 μM. Cell viability was assayed after five days using CellTiter-Glo 3D (Promega G9683), with luminescence measured. Drug response curves were generated by nonlinear regression using the percentage of viable cells against logarithm of drug concentration using Graphpad Prism 9. IC50 values were calculated by the equation log(inhibitor) versus response (variable slope, four parameters). Two-way ANOVA was used to compare dose-response curves. A similar method was used to determine drug response of PDX organoids and cell lines.

### ICB treatment of mice

Tumor-bearing mice were injected intraperitoneally every four days with either cocktail of anti-PD1 (250 μg/dose, #BE0146, BioXcell) and anti-CTLA4 (100 μg/dose, #BE0131, BioXcell) or control IgG (350ug/dose, #BE0089 and BE0087, BioXcell). Tumors were measured with calipers twice a week. On day 18, mice were euthanized, and tumors were collected for immunoprofiling.

### Immunoprofiling of T cells

Resected tumors were cut into small pieces using spring scissors and digested in 0.5 mg/mL collagenase D (Roche: cat#: COLLD-RO) and 0.25 mg/mL DNase I at 37°C for 30 minutes. After digestion, samples were passed through 70 μm cell strainers followed by ficoll density gradient centrifiguation (Lymphoprep: STEMCELL; cat# 07851). After removing erythrocytes, mononuclear cells were stimulated with phorbol 12-myristate 13-aetate (PMA), ionomycin, brefeldin A, and monensin in the T Cell-medium for four hours at 37°C. Cells were then blocked with anti-mouse CD16/32 (Biolegend; cat# 550994) at room temp for 1 minute, and stained with anti-CD90 (BioLegend; cat# 140327), anti-CD8 (BD Biosciences; cat# 560776), and anti-CD4 for 8 minutes in the dark. After staining, cells were washed and fixed/permeabilized using Perm-Fix buffer. Subsequently, cells were stained for anti-Ki67 (Thermo Fisher Scientific; cat# 56-5698-82), anti-TNFα (BioLegend; cat#506324), anti-IFN-γ (BD Biosciences; cat# 563773), and anti-Granzyme-B for 10 minutes. After further washing, the cells were analyzed on the BD LSRFortessa^TM^ Cell Analyzer, and flow cytometry data were analyzed using FlowJo V10.8.1.

### Drug treatment of mice

The anti-tumor efficacy of YJ9069 was evaluated in various subcutaneous xenografted and allografted models. In each case, when tumors reached ∼100-200 mm^3^, mice were randomized into two groups of 6-10 mice. Each group received either YJ9069 (15 mg/kg or 30 mg/kg) or vehicle (2 times/week) by IV injection for 14-30 days. Tumor volume was measured twice weekly by caliper following the formula (π/6)(LxW^2^) where L and W are the length and width of the tumors. At the end of the time course, tumors were excised, weighed, and collected for histological analysis.

### Immunoblotting

Cells were pelleted and lysed using 1X cell lysis buffer (Cell signaling, Cat# 9803S) with EDTA-free Protease Inhibitor Cocktail (Roche, Cat# 4693159001) and PhoSTOP (Roche, Cat# 04906837001). Protein concentration was determined using Pierce 660 nM Protein Assay Reagent (Thermo Fisher Scientific, Cat# 22660), and 20–30 μg of total protein was loaded in each lane. Proteins were separated by NuPAGE 3-8% or 4-12 % Tris-Acetate Midi Gel (Invitrogen, Cat# WG1402BX10) and transferred to nitrocellulose membranes (Fisher, Cat# 88018). Membranes were blocked with 5% non-fat dry milk /PBS for 1 h and then incubated with primary antibody overnight at 4°C. The primary antibody information is listed in **Table S2**. After three washes with 1 X TBS (ThermoFisher, Cat J75892-K8 pH7.4) containing 0.1% Tween-20 (Sigma, Cat P9416-100mL), membranes were incubated with 1:3000 diluted horseradish peroxidase (HRP) labeled secondary antibodies in 5% milk/PBS for 2 h at room temperature. After three washes with TBST, membranes were imaged using an Odyssey CLx Imager (LiCOR Biosciences). For the analysis of CDK12^as^ cells, anti-human antibodies were used for the following proteins: CDK12 (Cell Signaling 11973S, Abcam ab246887), β-Actin (Santa Cruz sc47778), α-TUBULIN (Santa Cruz 3873S), RNA polymerase II subunit B1 phosphor CTD Ser-2 Antibody, (clone 3E10, Millipore 04-1571). For CDK12^as^ cells, lysates were harvested with NP250 buffer (20 mmol/L Tris, pH 7.6, 1 mmol/L EDTA, 0.5% NP40, 250 mmol/L NaCl) containing protease inhibitor cocktail tablets (Roche). Samples were run alongside a Chameleon Duo protein ladder and transferred to nitrocellulose membranes, blocked using LICOR TBS blocking buffer (927-50000), developed using LICOR IRDye secondary antibodies, and imaged using an Odyssey CLx.

### CDK12^as^ survival assays

For siRNA experiments, cells were reverse transfected using Lipofectamine RNAimax Transfection Reagent (Promega) 6-well plates for 24 hours prior to splitting to final destination plates. For CDK12 cDNA experiments, cells were forward transfected with Lipofectamine 2000 Transfection Reagent in 6-well plates. After 24 hours, cells were divided into destination 6 well plates. 24 hours after seeding into destination 6-well plates, media containing small molecule inhibitors (talazoparib or 1NM) was added and replenished twice per week. After 2 weeks, colonies were washed with PBS, fixed with 10% trichloroacetic acid, and stained with sulforhodamine B. Image scans of stained colonies were analyzed for colony number and growth area by thresholding a grayscale image followed by conversion to a binary image with a watershed algorithm applied within ImageJ.

### CDK12^as^ γH2AX and Rad51 analysis

Cells were seeded in 96-well plates for 24 hours, exposed to indicated drug combinations for an additional 24 hours, or exposed to 10 Gy γ irradiation for 15 minutes. Cells were fixed with 4% PFA for 1 hour at room temperature and washed twice with PBS. Permeabilization was performed with 0.2% Triton x100 in PBS and blocked using a PBS solution with 1% BSA and 2% Fetal Bovine Serum. Primary antibody incubation (γH2AX using clone JBW301, Millipore 05-636 or RAD51 detection using Abcam ab63801) was carried out overnight at 4°C, and secondary antibody incubations were carried out for 40 minutes at room temperature. DAPI stain was added 10 minutes prior to development. Immunofluorescence was detected on an ImageXpress high content spinning disc microscope, and the number of foci per cell was determined with metaXpress software.

### siRNA screening and transfection

CDK12^as^ cells (1000 cells/well) were reverse transfected in a 96 well plate format with a custom siGENOME SMARTPool (Dharmacon) siRNA library as described previously^72^ using Lipofectamine RNAimax Transfection Reagent (Promega). Positive (siPLK1) and negative controls (siCON1, Dharmacon) were also included in each plate. After 24 hours, media was replaced with new media drug containing 1NM (0.3 μM) or the drug vehicle (DMSO), and cells were continuously cultured for six days further, at which point cell viability was estimated by the addition of CellTiter-glo reagent to the media for 10 minutes. Drug Effect Z scores were calculated from the resultant luminescence data as described previously^73^. Each screen was carried out in triplicate, with the data being combined in the final analysis. For single gene siRNA experiments, C4-2B cells were plated in 96-well plates and allowed to adhere overnight. The next day, cells were transfected using siGENOME SMARTPool (Dharmacon) against the indicated genes (CCNK, CDK13) or nontargeting control (NTC) as above. The plate was then placed in an IncuCyte S3 (Sartorius) and cell growth monitored over the indicated time frame.

### RNA-seq and data analysis

RNA extraction was followed by ribosomal RNA (rRNA) depletion. The rRNA-depleted RNA libraries were prepared using the KAPA RNA HyperPrep Kit (Roche) and subjected to the Agilent 2100 Bioanalyzer for quality and concentration. Transcripts were quantified by alignment-free approach kallisto^74^ using index generated from mouse reference genome (mm10) and then summed to obtain gene level counts. Differential analysis was performed using limma-voom procedure^75,76^ after TMM-normalization^77^ of gene level counts with calcNormFactors of edgeR^63^. Genes with mean Transcripts Per Million (TPM) less than 1 in both control and treatment groups were considered as lowly expressed genes and excluded for differential analysis. Enrichment of Hallmark gene sets downloaded from MSigDB^78^ were examined with fgsea^64^ using genes ranked by logFC estimated from limma as input.

### Whole-genome sequencing

Whole-genome sequencing was performed as per our standard protocols^18^. Briefly, tumor genomic DNA was purified using the AllPrep DNA/RNA/miRNA kit (Qiagen). *Cdk12^WT^* (reference genome) and *Cdk12^KO^* organoid-derived DNAs were sequenced on the Illumina NovaSeq 6000. Short reads were trimmed off sequencing adapters and aligned to the GRCm38 reference genome using BWA-MEM^79^, with settings "-Y -K 10000000", duplicates were removed per Picard^80^ rules, and depth of coverage was calculated using Mosdepth^81^ , with settings "-x -F 1796", excluding unmapped, not primary, QC-failed, and duplicate reads. Average depth of coverage in 10kb bins was normalized per sample to the total sequencing depth and adjusted for GC-bias using weighted LOWESS as implemented in Limma function loessfit^75^. The resulting coverage profiles were masked for outliers and segmented using CBS as implemented in DNACopy^82^. The resulting segmentation profiles were pruned using CNVEX as described previously^83^. The presence of focal gains was determined by identifying segments >50kb in size with a normalized log-coverage of > 0.5. The lack of FTDs was visually confirmed through visualizations of coverage profiles using R / ggplot2 and IGV.

### Generation of CRISPR knockout of *Cdk12*/*CDK12* in Myc-CaP cells and C4-2B

Short guide RNAs targeting the exons of mouse *Cdk12* were designed by Benchling (https://www.benchling.com/). Non-targeting control sgRNA and Cdk12-sgRNAs were cloned into lentiCRISPR v2 plasmid (Addgene 98290). Myc-CaP cells were transiently transfected with control sgNT or pair of two independent *Cdk12*-targeting sgRNAs. Twenty-four hours after transfection, cells were selected with 10 μg/mL puromycin for three days. Western blot was performed to detect knockout efficiency. Individual cells were isolated to generate monoclonal lines for analysis of knockout by Western blot. More than 100 clones were screened in this process. For C4-2B knockouts, cells were transfected with the PX458 plasmid (Addgene 48138) containing the guide sequence (CTTGGTATCGAAGCACAAGC or ACTTTGCAGCCGTCATCGGG) targeting exon 1 of *CDK12* using Lipofectamine 3000 (Thermo Fisher) according to manufacturer’s instructions. Approximately 48-72 hours after transfection, cells were sorted for green fluorescence protein (GFP) into single cells in a 96-well plate format. Clones were expanded and validated by Western blot and sequencing for the target site. Approximately 50 clones were screened in this process.

### Colony formation assay

Cells were seeded into six-well plates (1x10^4^ cells/ well) and allowed to grow for 5 days in complete medium. They were then fixed in 10% formalin for 30 minutes at RT and stained for 30 minutes in crystal violet (Fisher Chemical, C581-100) diluted to 1% by volume in H2O. Following H2O washes, samples were dried overnight and imaged on an Epson Perfection V33 scanner.

### Data and code availability

Sequencing data have been deposited in the National Center for Biotechnology Information Gene Expression Omnibus (NCBI GEO) with the accession number GSE254390. No custom code was developed in this study.

## SUPPLEMENTAL FIGURE LEGENDS

**Figure S1: *Cdk12* is partially ablated in prostate epithelium by *Probasin*-driven Cre recombinase. Related to Figure 1**.

**(A)** Prostate epithelial *Cdk12* ablation scheme.

**(B)** CDK12 immunohistochemistry (IHC) and *Cdk12* in situ hybridization (ISH) in 8-week-old WT and *Cdk12^pc-/-^* mice.

**(C)** Percent epithelial cells immunonegative for CDK12 (*Cdk12* KO cells) in prostate lobes of *Cdk12^pc-/-^* mice: anterior prostate (AP), ventral prostate (VP), dorsal prostate (DP), lateral prostate (LP). (n= 2-3 prostate cross sections from 6 mice).

**Figure S2: C*dk12* ablation in prostate epithelium of mixed background mice causes pre-cancerous lesions with aging. Related to Figure 1**.

**(A)** Hyperplasia with lost nuclear polarity and isonucleosis in prostate epithelium of 30-week-old mixed background *Cdk12^pc-/-^* mice. Note concentrated Ki67 staining in histologically abnormal regions. These regions are absent in wild-type (WT) controls.

**(B)** Larger pre-cancerous lesions (indicated by dashed line) in prostate epithelia of 52-week-old mixed background *Cdk12^pc-/-^* mice.

**(C)** Percent cross sectional area occupied by pre-cancerous lesions in prostate lobes of 52-week-old *Cdk12^pc-/-^* mice. Anterior prostate (AP), ventral prostate (VP), dorsal prostate (DP), lateral prostate (LP). (n= 2-3 prostate cross sections from each of 6-7 mice). *p<0.05, **p<0.01.

**Figure S3: Application of *mT/mG* model to isolate cells with active Cre recombinase and *Cdk12* ablation; demonstration of abnormal morphology in organoids generated from *Cdk12*-null cells. Related to Figure 2**.

**(A)** Generation of a *Pb-Cre*;*Cdk12^f/f^*;*mT/mG* prostate mouse model to identify prostate epithelial cells with active Cre recombinase.

**(B)** Basal cell isolation from *Pb-Cre*;*Cdk12^f/f^*;*mT/mG* prostate (52-week time point).

**(C)** *Cdk12* mRNA expression in 52-week *Pb-Cre*;*Cdk12^f/f^*;*mT/mG* prostate epithelial cells. BC, basal cells; LEC, luminal epithelial cells.

**(D)** Organoids derived from *Pb-Cre*;*Cdk12^f/f^*;*mT/mG* prostate basal cells (52-week time point): Red/ Tom(+) contain intact, wild-type *Cdk12* (*Cdk12^WT^*); green/ GFP(+) have *Cdk12* ablation (*Cdk12^KO^*).

**(E)** Confirmatory experiment demonstrating that EGFP(+)/ Cre-expressing cells from *Cdk12^+/+^* mouse prostate do not have abnormal organoid phenotype.

**(F)** Acute *Cdk12* ablation achieved through *in vitro* adenoviral Cre to *Cdk12^f/f^*;*mT/mG* organoids. Ad-CTRL indicates control adenovirus. Ad-Cre indicates Cre-expressing adenovirus. Ablation of *Cdk12* gene is coupled with red (Tom) to green (GFP) color change. Images show morphology of organoids with WT *Cdk12* (Ad-CRTL) and *Cdk12* ablation (Ad-Cre).

**Figure S4: *Cdk12* null mice exhibit upregulation of specific prostate cancer-associated pathways and p53 target genes. Related to Figure 3**.

**(A)** Scheme for scRNA-seq analysis of prostates from *mT/mG* mice with wild-type (WT) Cdk12 (*Cdk12^+/+^*) or prostate epithelial-specific *Cdk12* ablation (*Cdk12^f/f^*) driven by *Probasin*-Cre (*Pb-Cre*) (KO). Cells of mT/mG mice express td-Tomato (TdTom) at baseline. The TdTom sequence is excised in cells with active Cre recombinase, enabling expression of enhanced GFP (EGFP). The strategy outlined above allows for comparison of only Cre-expressing cells (i.e., EGFP-expressing cells) from each animal. (n= 3 mice per group)

**(B)** UMAP plot indicating cell populations from cells described in (A). Cells are annotated using the Crowely et al. dataset as reference^60^. LumD, LumL, and LumV are Luminal cells specific to dorsal, lateral, and ventral prostate, respectively; LumP is the proximal progenitor population.

**(C)** Enrichment of Lum P population in *Cdk12*-null GFP(+) cells (*Cdk12^KO^*).

**(D)** Enrichment of cancer-related pathways in LumV cells of the *Cdk12^KO^* prostate epithelium.

**(E)** Enrichment plots of selected pathways enriched in luminal cells from the *Cdk12^KO^* ventral prostate.

**Figure S5: Clonal *Cdk12^KO^*organoid lines do not demonstrate focal tandem duplications. Related to Figure 4**.

**(A)** *Cdk12/Trp53* double KO organoid cells serially passaged as subcutaneous allografts in mice.

Each line represents an individual allograft.

**(B)** Immunohistochemical staining of CDK12, γH2AX, AR, Krt8, and p53 in *Cdk12^KO^*-sgp53 allografts, Myc-CaP allografts, and TRAMP-C2 allografts, and prostates of the established *Pten^pc-/-^* prostate cancer mouse model. Scale bar indicates 50μm

**(C)** Genomic sequencing of clonal *Cdk12^KO^* organoid lines (*Cdk12^KO^*-1, *Cdk12^KO^* -2, *Cdk12^KO^*-3, *Cdk12^KO^*-4) demonstrating ablation of exons 3 and 4. Each plot indicates sequencing of an individual monoclonal organoid line.

**(D)** Genomic sequencing of *Cdk12^KO^* organoid lines demonstrating that these do not have evidence of the focal tandem duplication pattern seen in human prostate cancer lacking functional *CDK12*. Each plot indicates sequencing of an individual monoclonal organoid line.

**Figure S6: Validation of CDK12^as^ cells and overview of siRNA screen. Related to Figure 6**.

**(A)** Diagram indicating generation of an analog sensitive CDK12 mutant HeLa cell line (CDK12^as^) (in lab of Arno Greenleaf) by two rounds of CRISPR mutagenesis; the first introduced a stop mutation into one *CDK12* allele and the second introduced a homologous recombination template coding for an p.F813G amino acid change in the DFG sequence of the CDK12 kinase domain. This alteration causes the kinase domain to take a wider configuration and to be inhibited by the adenine analog 1-NM-PP1 (1NM).

**(B)** Western blot indicating the effect of CDK12 inhibition on phosphorylation of the CTD of RNA Pol-II in CDK12^as^ cells exposed to 1NM.

**(C)** Quantification of relative protein level shown in (B) from three independent experiments. Error bars represent standard error of the mean (SEM). p value calculated by t-test.

**(D)** Colony formation assay (CFA) images from CDK12^as^ or CDK12 wild type cells (CDK12 ^wild^ ^type^) exposed to 1NM and/or the PARP inhibitor talazoparib.

**(E)** Quantification of CFA data from replica experiments. Dots represent the mean and error bars represent SEM. p value calculated by two-way ANOVA.

**(F, G)** Quantification of irradiation-induced nuclear γH2AX and RAD51 nuclear foci in CDK12^as^ cells exposed to 1NM. Dots represent the percentage of cells with 5 or more detectable nuclear foci. Columns indicate the median score from five experiments. Error bars represent SEM. p values calculated by t-test.

**(H)** Western blot showing overexpression of a wild-type CDK12 cDNA construct in CDK12^as^ cells.

**(I)** Wild-type CDK12 expression reduces 1NM-induced PARP inhibitor sensitivity. CellTiter-Glo results from 1 week of growth normalized to DMSO treated controls. Columns indicate the median of 6 biological replicate samples, indicated by individual dots. p value calculated by t-test.

**(J)** Schematic of siRNA screen. CDK12^as^ cells were transfected in a 96 well plate format with a custom siRNA library targeting genes annotated as shown. Positive (siPLK1) and negative controls (siCON1, Dharmacon) were also included in each plate. After 24 hours, media was replaced with new media drug containing 1NM (0.3 μM) or the drug vehicle (DMSO) and cells were continuously cultured for six further days, at which point cell viability was estimated by the use of CellTiter-glo reagent.

**(K, L)** Quality control data from siRNA screen. Normalized percent inhibition (NPI) data is shown for non-targeting control siRNA (siCON1, normalized at NPI = 1 from three replica screens), siRNA targeting PLK1 (normalized at NPI = 1 from three replica screens), and siRNA designed to target each of the genes included in the screen. Data from DMSO-exposed (K) and 1NM-exposed (L) arms of the screen are shown, indicating a large dynamic range between siCON1 and siPLK1.

**Figure S7: CDK13/12 inhibitor treatment in organoids. Related to Figure 6**.

(A) *Cdk12^WT^*-sgp53 and *Cdk12^KO^*-sgp53 organoids treated with CDK13/12 inhibitors (YJ5118, THZ531). Bright field images show organoids subjected to each agent at concentrations of 200 and 500 μm (and DMSO vehicle control). Line graphs show IC50 curves for both organoid types treated with each agent. (n= 3 samples per group, 2 individual experiments).

(B) C4-2B *CDK12*KO and C4-2B CTRL cells: Colony formation assay performed with increasing THZ531 concentrations.

(C) C4-2B *CDK12*KO and C4-2B CTRL cells: Percent viability at increasing THZ531 concentrations.

(D) IC50 values of *Cdk12* knockout (sgCdk12) and control (sgNT) Myc-CaP cells treated with CDK12/13 degrader YJ9069.

**Figure S8: PDX model with *CDK12* loss of function. Related to Figures 6-7**.

**(A)** CDK12 IHC of the indicated prostate PDX lines. Scale indicates 50μm.

**(B)** PDX line LTL706B (biallelic frameshift mutations in *CDK12* gene) in mouse kidney. Tumor tissue stained for AR, KRT8, PSMA, and CDK12. Scale indicates 50μm.

**(C)** Scheme for organoid generation from LTL706B tumors. Organoids derived from LTL706B tumors in bright field (left panel, scale 1000μm; right panel, scale 200μm) and embedded/cross-section stained for AR, KRT8, PSMA, and CDK12 (scale 50μm).

**(D-E)** Unmodified (sgNT-treated) Myc-CaP allografts (E) and sgCdk12-treated Myc-CaP allografts (F) subjected to *in vivo* treatment with the orally bioavailable CDK13/12 degrader YJ1206. Line graphs indicate tumor volumes, and bar graphs indicate tumor weight at end of treatment time course. (n= 7-8 mice, with 2 tumors per mouse, per group).

## SUPPLEMENTAL TABLES

**Table S1:** siRNAs used for synthetic lethality screening in CDK12^as^ cells.

**Table S2:** sgRNAs, Taqman probes, QPCR primers, and antibodies.

