## Supplementary figures and images for "*CDK12* Loss Promotes Prostate Cancer Development While Exposing Vulnerabilities to Paralog-Based Synthetic Lethality"

### Supplemental Figures S1 - S8

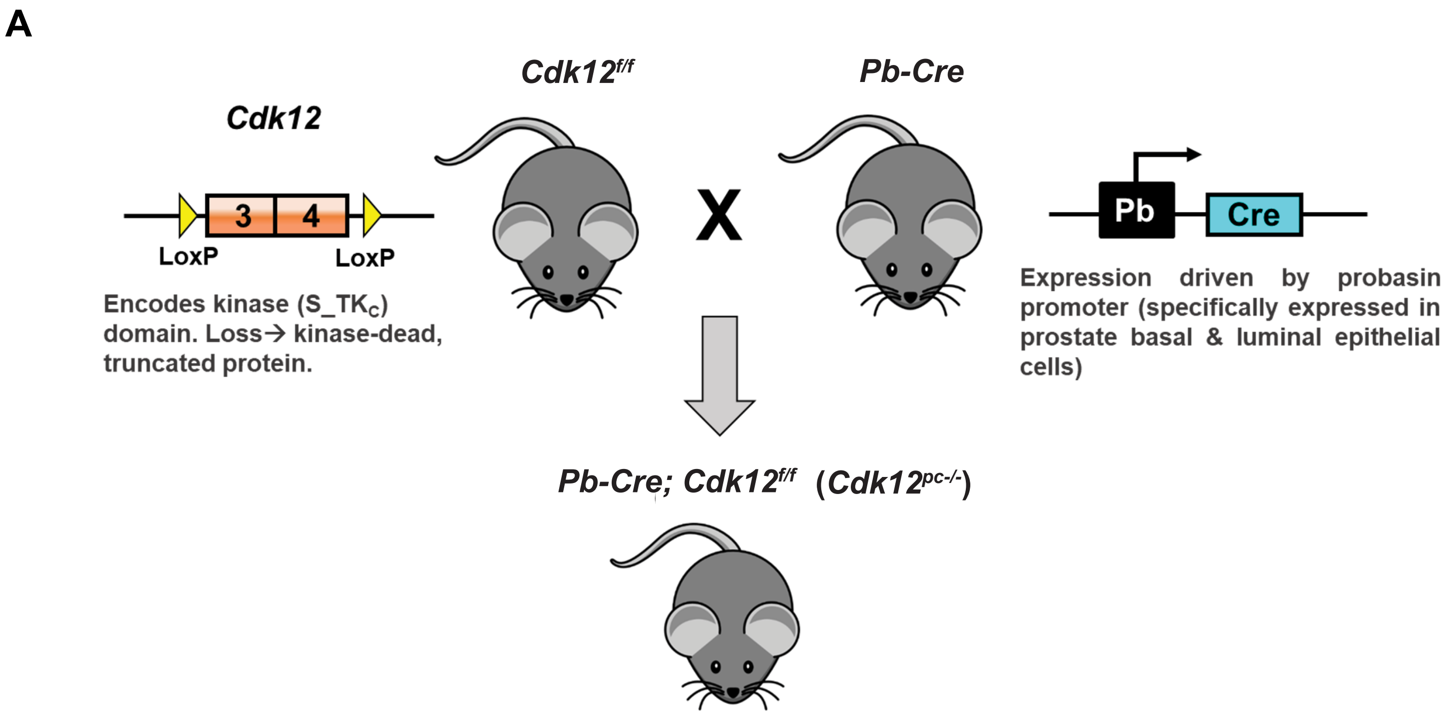

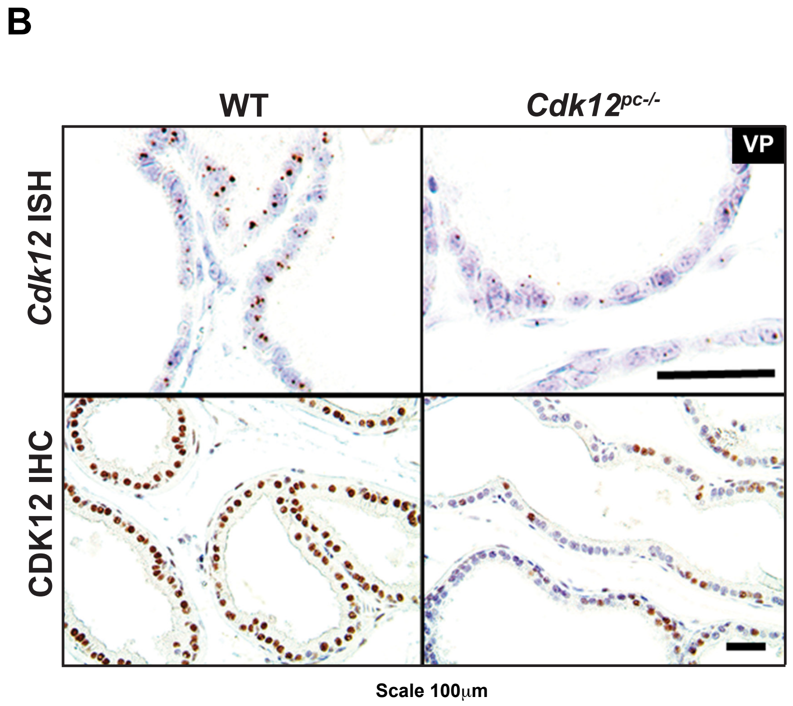

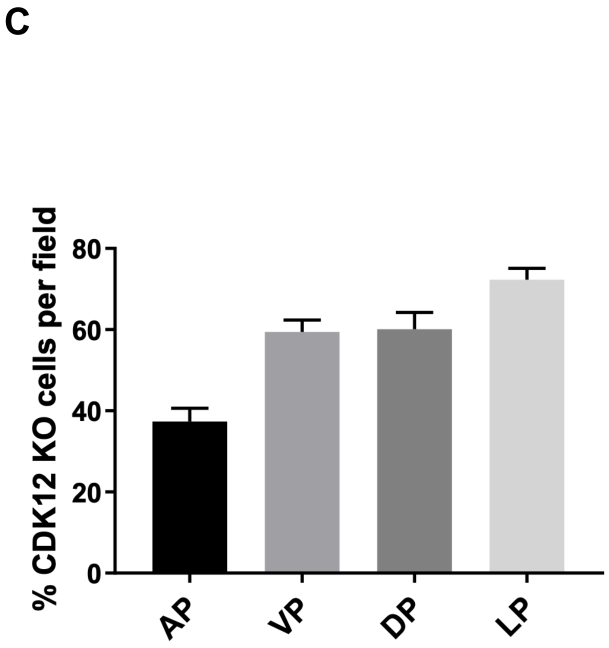

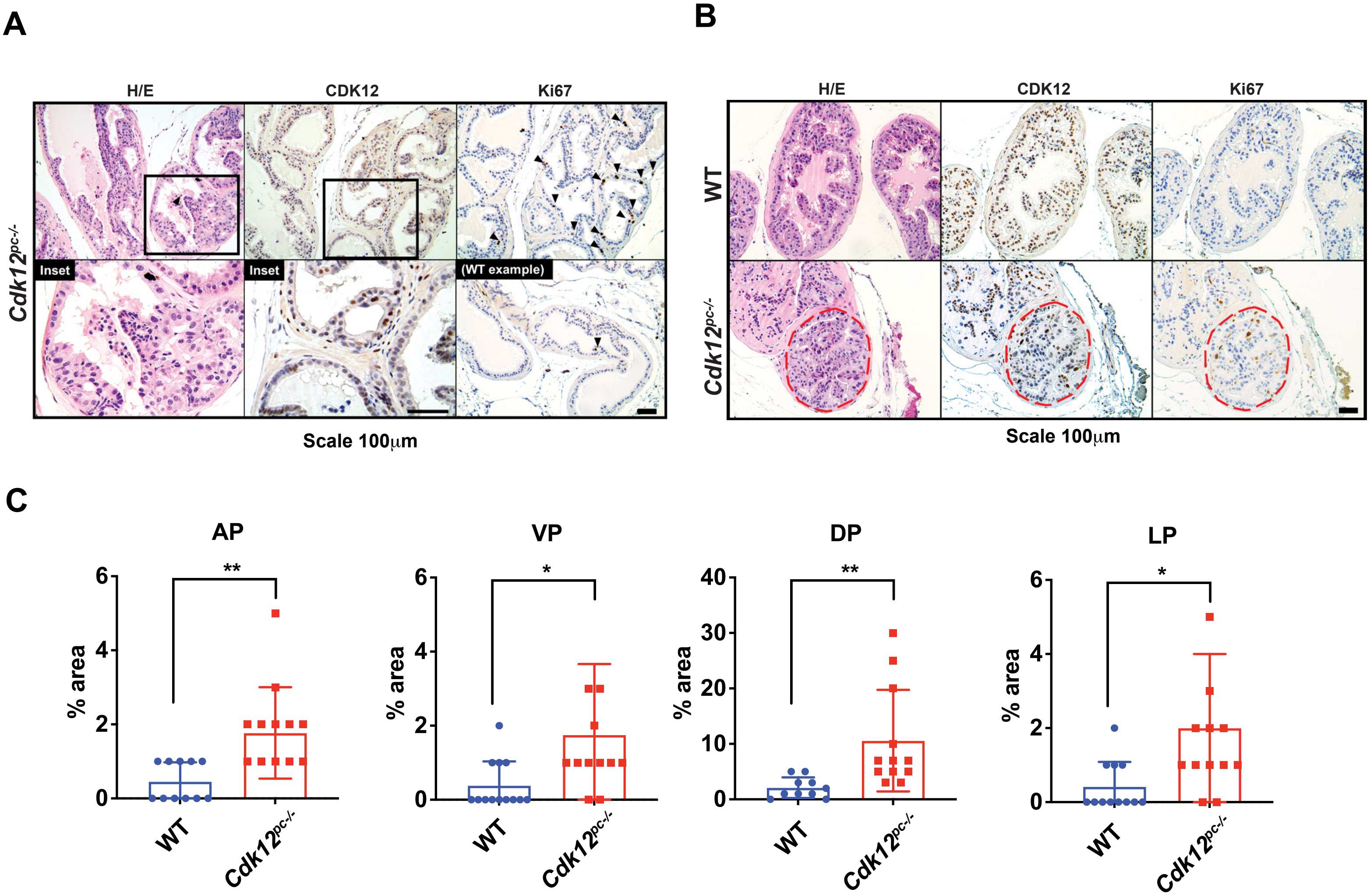

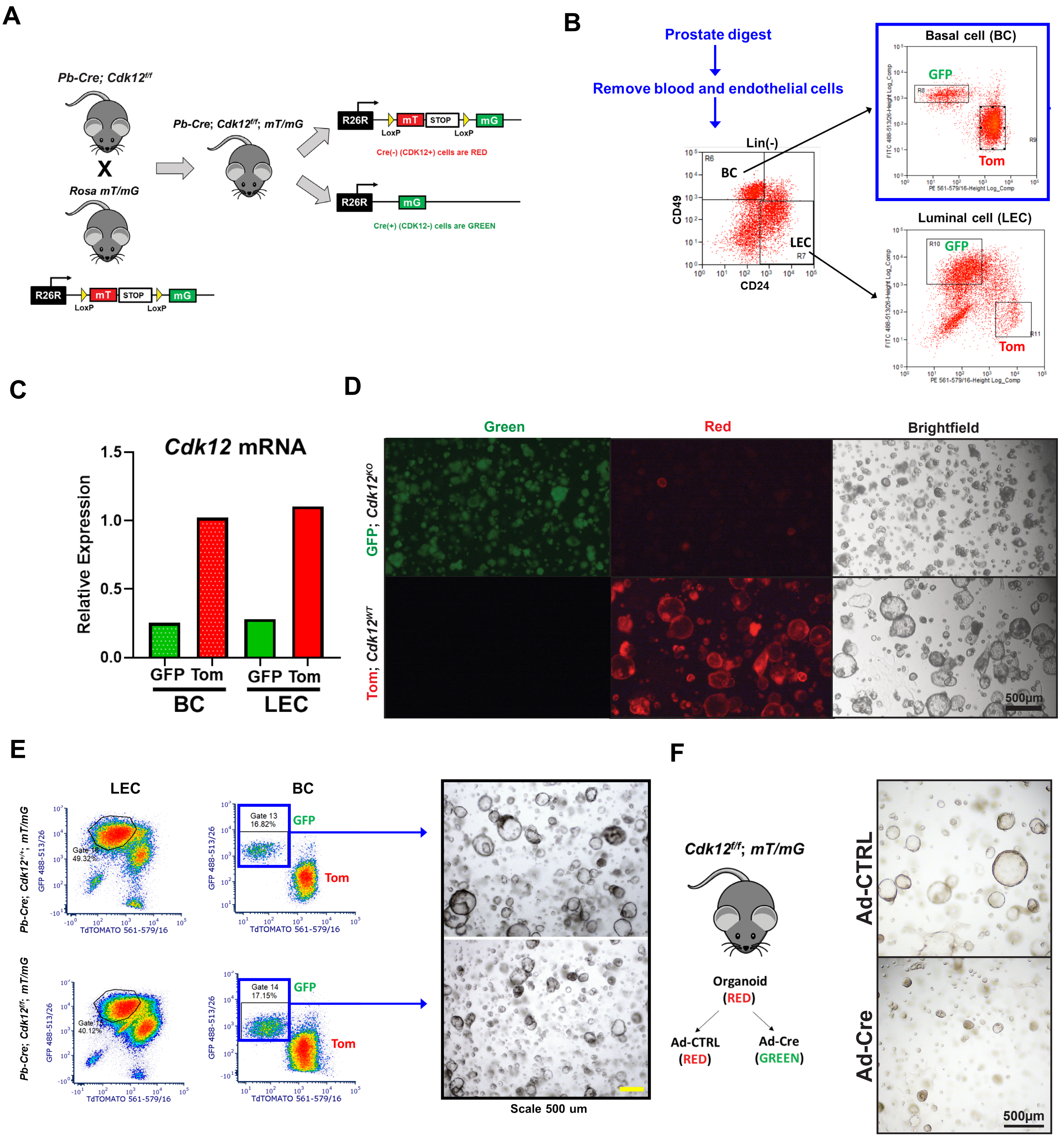

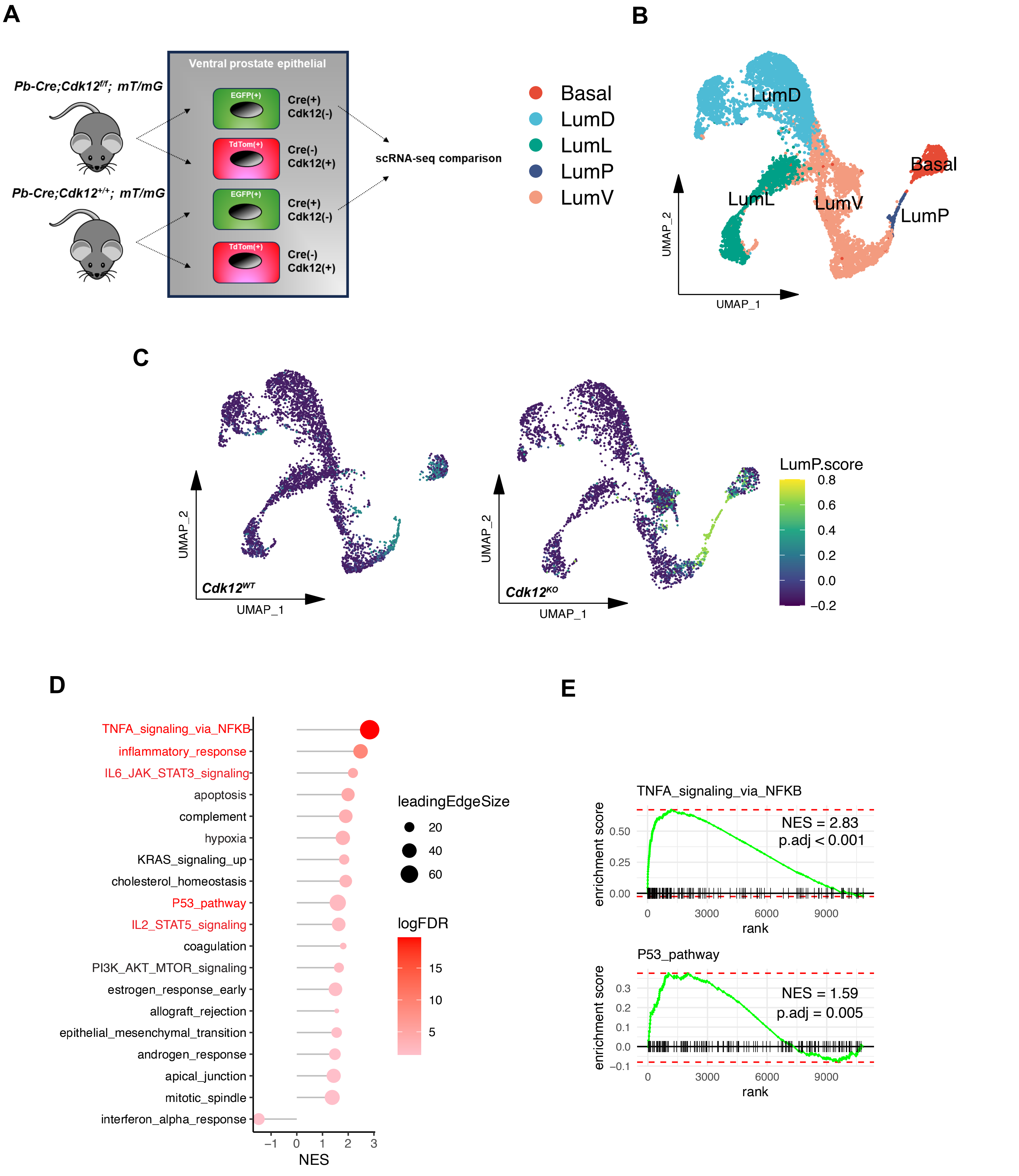

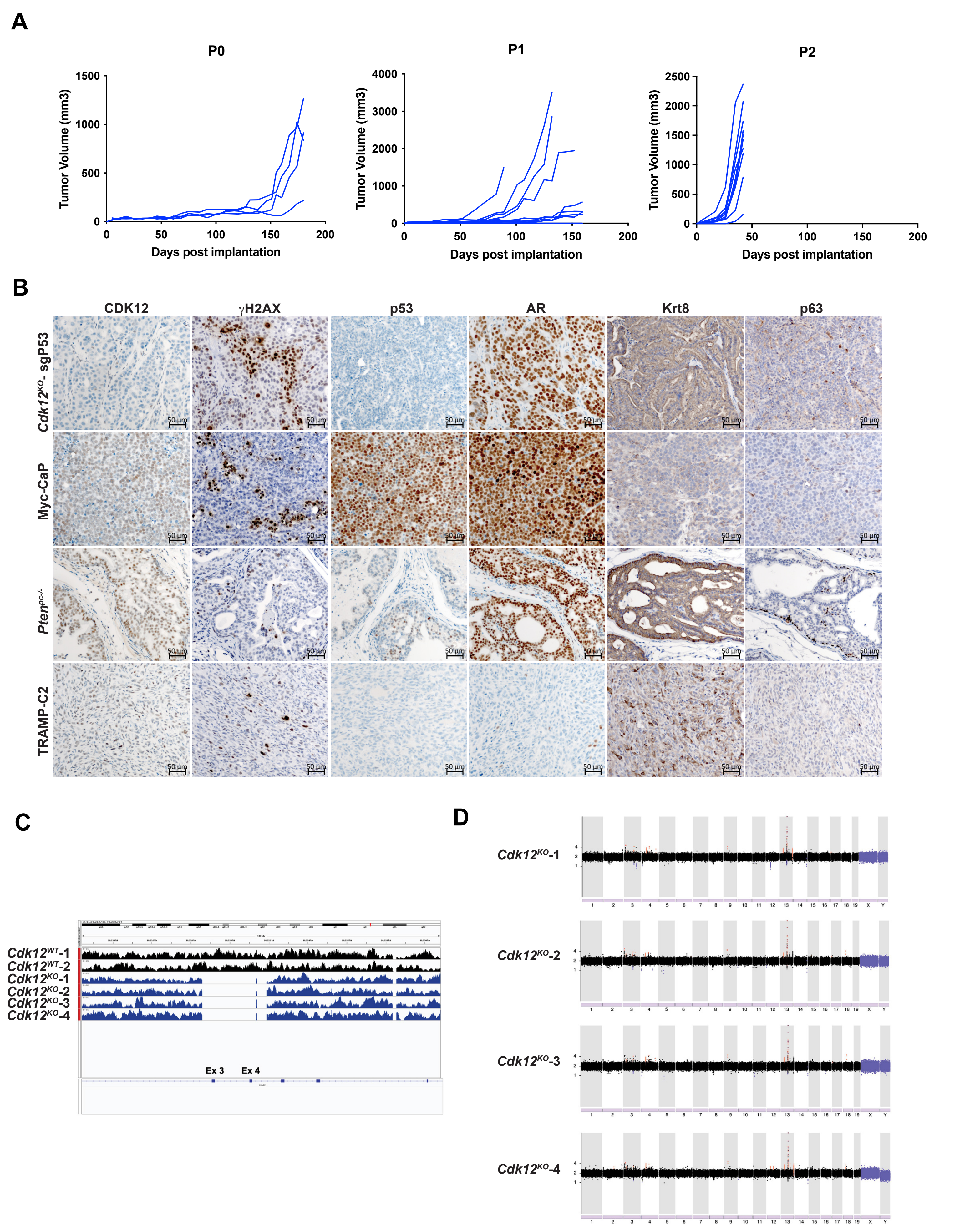

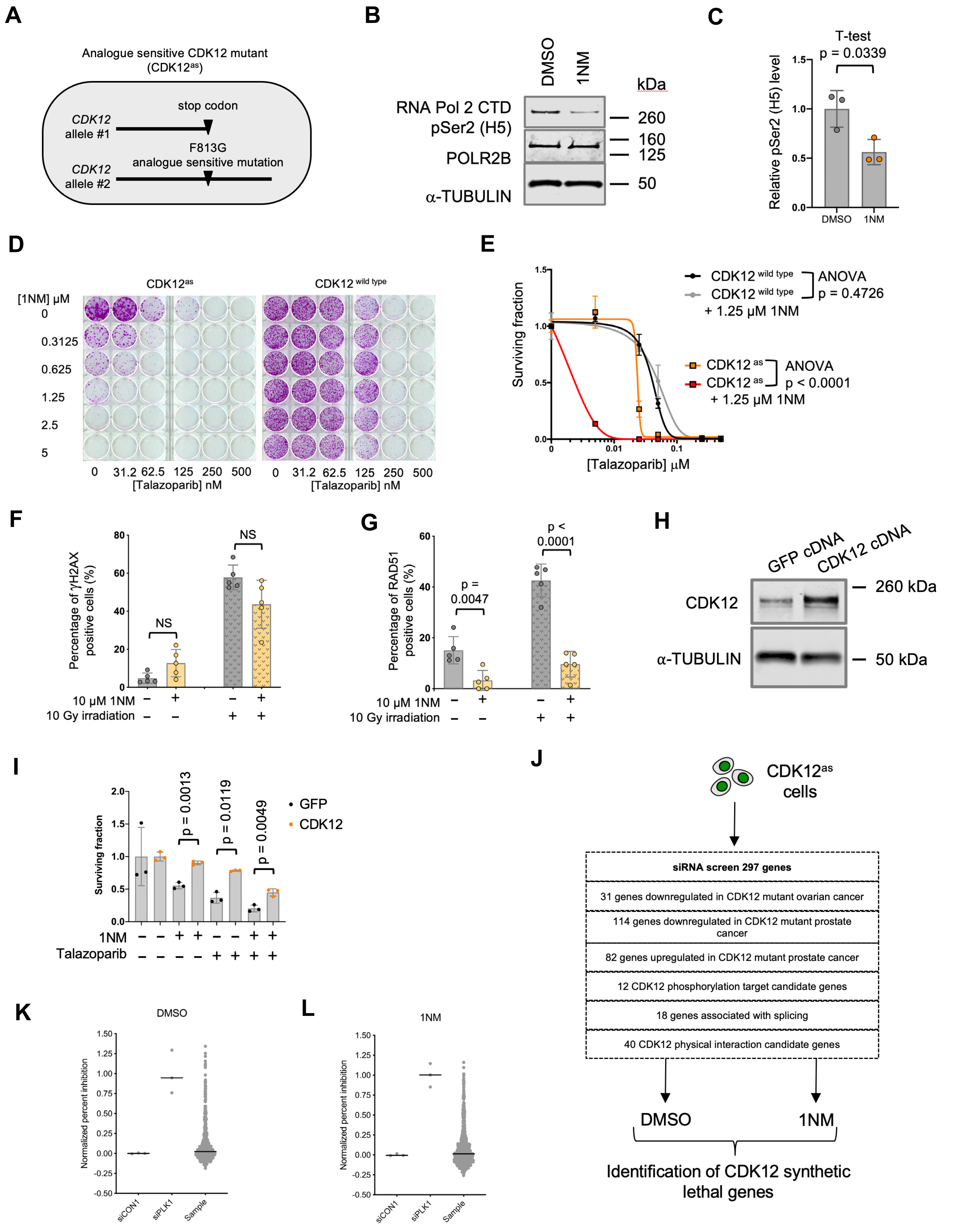

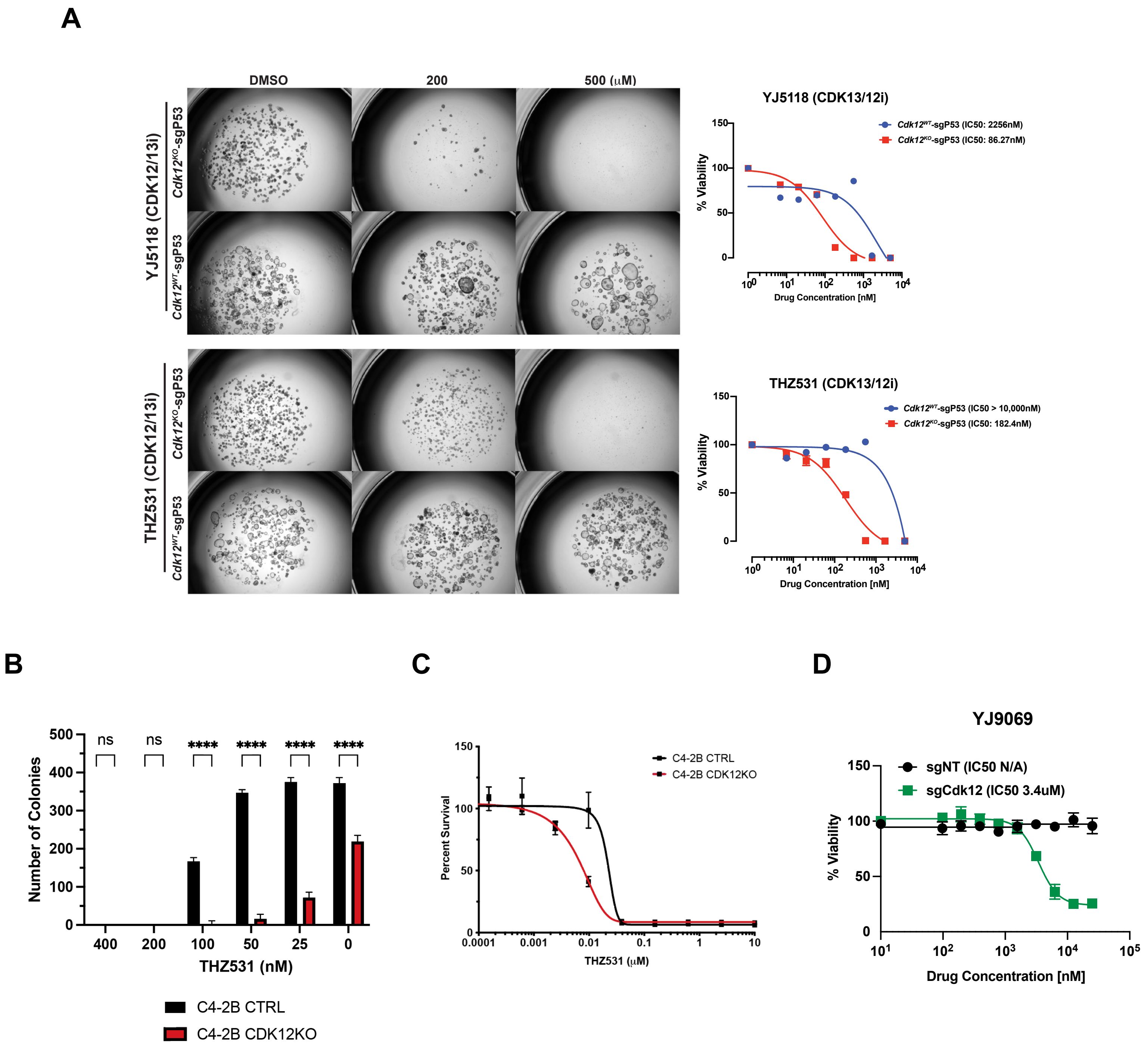

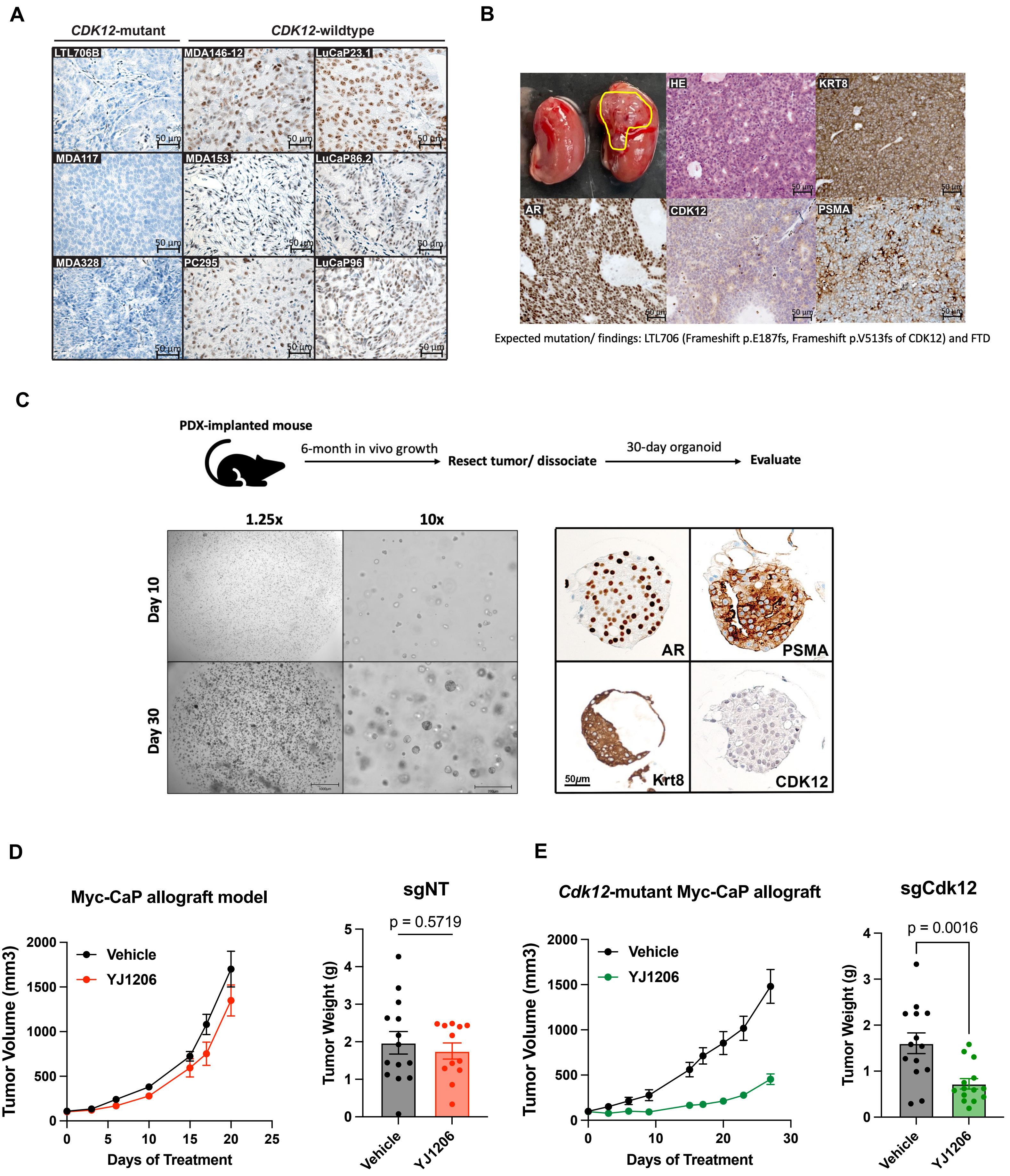
